# Single-molecule tracking-based drug screening

**DOI:** 10.1101/2023.11.12.566743

**Authors:** Daisuke Watanabe, Michio Hiroshima, Masato Yasui, Masahiro Ueda

## Abstract

The single-molecule tracking of transmembrane receptors in living cells has provided significant insights into signaling mechanisms, such as mobility and clustering upon their activation/inactivation, making it a potential screening method for drug discovery. Here we show that single-molecule tracking-based screening can be used to explore compounds both detectable and undetectable by conventional methods for disease-related receptors. Using an automated system for a fast large-scale single-molecule analysis, we screened for epidermal growth factor receptor (EGFR) from 1,134 of FDA approved drugs. The 18 hit compounds included all EGFR-targeted tyrosine kinase inhibitors (TKIs) in the library that suppressed any phosphorylation-dependent mobility shift of EGFR, proving the concept of this approach. The remaining hit compounds are not reported as EGFR-targeted drugs and did not inhibit EGF-induced EGFR phosphorylation. These non-TKI compounds affected the mobility and/or clustering of EGFR without EGF and induced EGFR internalization, to impede EGFR-dependent cell growth. Thus, single-molecule tracking provides a new modality for discovering novel therapeutics on various receptor functions with previously untargeted mechanisms.

## Introduction

Transmembrane receptors trigger signal transduction to induce cellular responses. Since the downstream signaling depends on the functional properties of receptors, various diseases are often attributed to receptor dysfunction. In drug discovery, the largest number of target molecules are membrane receptors (> 50% of total), such as receptor tyrosine kinases (RTKs), G protein-coupled receptors (GPCRs), and immune receptors, followed by nuclear receptors (< 20%)^1–3^. Epidermal growth factor receptor (EGFR), an RTK, mediates signal transduction for cell proliferation, differentiation, migration, and apoptosis, and is a primary target molecule in drug exploration because its overexpression and/or mutations are found in various types of cancer cells^4–7^. Many small-molecule drugs against a well-known EGFR-related cancer, non-small cell lung cancer (NSCLC), have been developed: gefitinib, erlotinib, and icotinib as first-generation tyrosine kinase inhibitors (TKIs); afatinib and dacomitinib as second-generation TKIs; and osimertinib as third-generation^8^. These drugs have been evaluated by their inhibitory effects on tyrosine kinase activity^9^, but EGFR undergoes multiple signaling processes including tyrosine phosphorylation, oligomerization, coupling with adaptor molecules, and internalization on the membrane of living cells^10^. Disturbing these processes can affect the signaling activity of the molecule, providing a potential mechanistic target for drug discovery. In general, assays for drug discovery focus on a particular step in the signaling pathway exhibited by the targeted molecules. However, assays that select for compounds that have effects on multiple steps in the signaling pathway are of interest because of their different mechanisms of action on the same target^11^.

Single-molecule imaging analysis has been used to visualize and measure the location and brightness of individual membrane receptor molecules in cells^12–14^. These measurements provide information on the lateral diffusion and oligomerization/clustering of receptors on the membrane of living cells. In the case of EGFR, the behavioral transition on the membrane, which is described by EGFR mobility and clustering dynamics, has been shown to correlate to the signaling process^15–19^: EGF triggers tyrosine phosphorylation, leading to the translocation of EGFR along the membrane to a specific region where the mobility of EGFR is confined and the clusters responsible for downstream signaling are formed, followed by internalization into endosomes. Single-molecule imaging has also revealed other processes during EGFR signaling, such as ligand-receptor binding kinetics, molecular structure transitions depending on membrane components, and colocalization with receptors for different signaling pathways. That is, single-molecule imaging has the potential to elucidate the effects of pharmacological compounds on multiple events during a series of EGFR signaling processes. In particular, this technique can detect changes in the physical properties of the target receptor, which include the lateral diffusion and the statistical distribution of receptor clusters, both of which are difficult to detect using biochemical methods. However, the low throughput of single-molecule imaging for data acquisition, which arises from its expertise- and manual-dependent workflow, prevents this method from being employed for large-scale single-molecule analysis. As a solution, we previously achieved a fully automated, in-cell single-molecule imaging system equipped with robotics and machine learning (AiSIS)^20,21^ that has a 100-fold higher throughput than standard single-molecule analysis techniques. High-throughput analysis by AiSIS enables the evaluation of numerous compounds by detecting their effect on molecular mobility and clustering during the signaling process, which provides a form of molecular-targeted drug screening.

Here, we report a single-molecule tracking-based technique for the screening of compounds targeting transmembrane receptors. To prove the concept of this approach, we screened for EGFR from a library consisting of 1,134 FDA approved compounds and successfully selected all TKIs contained in the library. The other hit compounds exhibited effects on the clustering, internalization, and expression level of EGFR without strong inhibition of the tyrosine kinase activity upon EGF stimulation. Different from screening methods that focus on a single event in the molecular process, our method could assess various physical events at the single-molecule level occurring during the signal transduction of the target molecule.

## Results

### EGFR mobility-based screening

AiSIS allows for a large-scale, single-molecule analysis of plasma membrane receptors using 96-well plates (Fig. 1a). Cells in each well were treated with a compound, and single-molecule imaging using total internal reflection fluorescence microscopy (TIRFM) was automatically executed well by well. Novel autofocusing (Fig. 1b and Supplementary Fig. 1) kept the in-focus position precise to acquire high magnification images of living cells with clear single fluorescence spots of the target molecules on the membrane by TIRFM. The fields of view suitable for the single-molecule analysis were determined by machine learning-based automated cell searching. During the imaging, autofocusing was continuously running, and the ligand was dispensed at desired times by robotics. In the case of measuring 20 cells under one measurement condition, as described later, 480 different measurement conditions for ligand/compound concentrations, time of the addition, and duration of the treatment could be examined on demand in a single assay in one day. By observing individual cells under uniform fluorescence excitation at high magnification, quantitative analysis of the position as well as the fluorescence intensity of the fluorescent spots is possible, which allows the characterization of not only the lateral mobility of the target molecule but also the cluster formation of the target molecule. Fig. 1c shows a single EGFR-mEGFP complex expressed in CHO-K1 cells, which lack endogenous EGFR. From the images acquired under optimized conditions, trajectory data, including the positions and brightness of fluorescent EGFR, were obtained by single-molecule tracking (Fig. 1d) and used to quantify the mobility of EGFR with diffusion coefficients and mean square displacements (MSD) (Fig. 1e). The mobility decreased with EGF addition, as shown by the MSD (Fig. 1f), consistent with a previous study^20^. This EGF-induced mobility shift was recovered by treatment with a TKI (gefitinib; Fig. 1e, bottom), which confirmed EGFR mobility tightly correlated with EGF-activated kinase function. As shown in Fig. 1f, a linear relationship was found between the MSD for 500 ms (MSD_500ms_) and EGFR phosphorylation under various EGF concentrations. Both data were obtained approximately 2 min after EGF addition, a time that represents the early phase of EGFR signaling just after reaching maximum EGFR phosphorylation. The inhibitory effect of gefitinib on EGF-induced decrease of MSD_500ms_ is shown in Fig. 1g (top). Because TKI itself seems to have little effect on the mobility in our data, the MSD ratio (MSD with EGF to MSD without EGF) was used to evaluate the compounds en bloc. As shown in Fig. 1g (bottom), when the EGF concentration was 30 nM, the ratio was approximately 0.2 for TKI-untreated cells but approached 1 for treated cells due to suppression of the EGF-induced decrease of EGFR mobility.

**Fig. 1.**
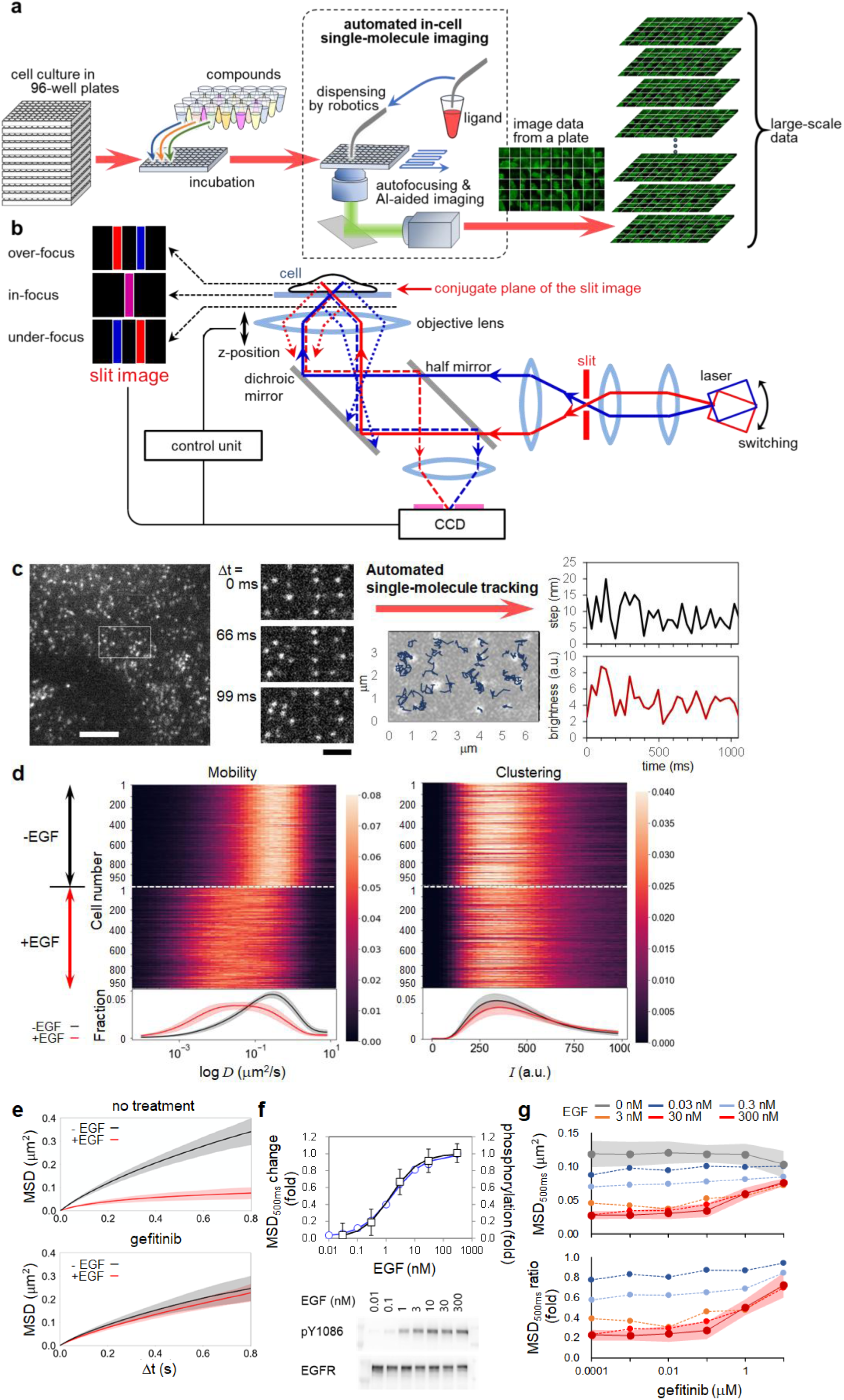
Large-scale single-molecule imaging analysis of fluorescently labeled EGFR for drug screening. **a,** Schematic flow of single-molecule imaging using AiSIS for drug screening. **b,** Optics of the autofocus device. Slit images, which reflect deviation from the in-focus position of the objective lens, are generated with switching illumination and used for feedback control. **c,** Automated single-molecule tracking. Left, representative images of EGFR-mEGFP observed with AiSIS. Scale bar, 5 μm. Middle left, time series images of the squared region in the left panel. Scale bar, 2 μm. Middle right, trajectories of EGFR (> 8 frames). Right, time series of step length (top) and brightness (bottom) of a spot in the left panels. **d,** Heatmaps of diffusion coefficients and brightness measured before and 5 min after the EGF addition. The average distributions over the cells are shown at the bottom. Black and red lines with shaded areas indicate the average with SD before and after EGF addition, respectively. **e,** Time development of MSDs for either untreated or gefitinib-treated (10 μM for 1 hour) cells without (black) and with EGF (300 nM, red). The acquisition of single-molecule images began 1 minute after the EGF addition. **f,** Relative changes in MSD_500ms_ and fold changes of phosphorylation obtained from western blotting (bottom) against different EGF concentrations. Single-molecule imaging and the phosphorylation assay began 1 and 2 minutes after the EGF addition, respectively. **g,** Dose-dependency of gefitinib on MSD. Top, MSD versus gefitinib concentration. Bottom, ratios between MSD with and without EGF. Colored curves correspond to the indicated EGF concentrations. Single-molecule imaging began 1 minute after the EGF addition. Shaded areas (c, d, and f) and error bars (e) indicate the SD and SE, respectively.

We extended the single-molecule tracking-based assay to drug screening. First, we optimized the conditions of the image acquisition (Supplementary Fig. 2), and the suitability of this approach as a screening method was assessed using the indices calculated from MSD_500ms_, with the positive control being cells treated with gefitinib and EGF and the negative control being cells treated only with EGF (Fig. 2a). The coefficient of variation (CV), signal/background (S/B), and Z’-factor were respectively defined as the ratio of the standard deviation (SD) and average of the controls, the ratio between the averages of the positive (signal) and negative (background) controls, and the following equation:

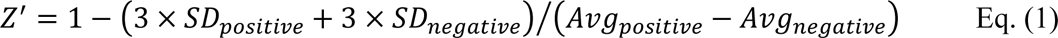

**Fig. 2.**
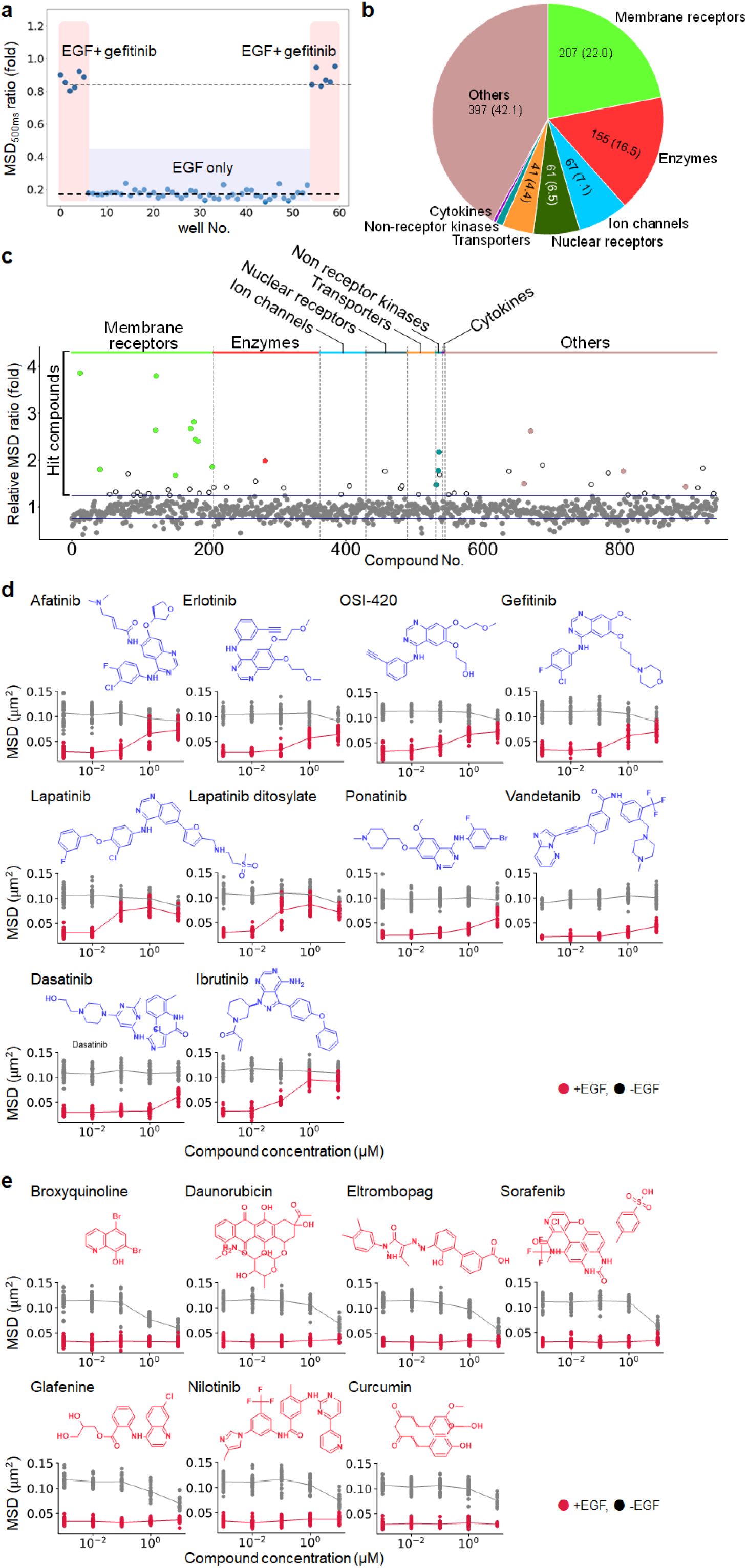
EGFR mobility-based screening for compounds in an FDA-approved library. **a,** Evaluation of single-molecule tracking as a screening method. MSD_500ms_ ratios of EGFR were calculated with and without gefitinib as positive and negative controls, respectively. Red-shaded regions indicate the MSD_500ms_ ratio obtained from wells treated with 30 nM EGF and gefitinib (positive control), and the blue-shaded region indicates wells treated with only EGF (negative control). Single-molecule imaging began 1 minute after EGF addition. **b,** Target proteins of FDA-approved compounds in the library. **c,** Effects of compounds on EGFR mobility were evaluated using the MSD_Δt_ ratio in which Δt provided optimal duration to obtain the best Z’-factor value in each well plate (Supplementary Fig. 3a). Ratios for compound-treated cells were normalized to that for untreated cells. Blue lines denote upper and lower thresholds. The 53 hit compounds beyond the upper threshold are indicated by the colored and white circles. After further evaluation, the selected 18 compounds were denoted by colored circles, in which the colors indicate the type of target protein. **d** and **e,** The dose-dependency was further evaluated by referring MSD_167ms_ against a series of compound concentrations. Red and black circles denote MSD values with and without EGF, respectively. 10 compounds (except auranofin) suppressed the EGF-dependent decrease in MSD (d), and 7 compounds reduced MSD regardless of the EGF concentration (e).

, where *SD***_positiv_**_e_, *SD_negative_*, *Avg***_positiv_**_e_, and *Avg_negative_* represent the SD and average for the positive and negative controls, respectively. The obtained indices for our system were 4% for CV, 2.7% for S/B, and 0.64 for Z’-factor. All these values were shown to satisfy the thresholds of ≤ 10% for CV, ≥ 2% for SB, and ≥ 0.50 for Z’-factor for drug screening (Supplementary Table 1). Next, we carried out the screening using a library of 1,134 FDA-approved compounds whose target proteins are responsible for broad functions such as receptors, channels, kinases, enzymes, transporters, and so on (Fig. 2b and Supplementary Table 3). This library includes 7 marketed drugs that act as TKI for EGFR (afatinib, erlotinib, OSI-420 (the active metabolite of erlotinib), gefitinib, lapatinib, lapatinib ditosylate (another form of lapatinib), and vandetanib). To explore compounds significantly affecting EGFR mobility, the screening procedure was designed as follows: 1) cultured cells in each well of a 96-well plate were pretreated with 100 μL of 10 μM compound solution at 37°C for 1 hour, 2) single-molecule imaging was automatically done on 20 different cells, 3) 100 μL of 120 nM EGF solution was added by the automatic dispenser so that the final concentration of EGF was 60 nM, and 4) another 20 cells were imaged 2 min after the EGF addition. During the screening, the Z’-factor was calculated for every assay plate from positive- and negative-control wells (20 cells in each well) to evaluate the screening quality, and images acquired from the plates passing the quality inspection were used in the following analysis (Supplementary Fig. 3a). To achieve a highly accurate screening, MSD_Δt_ was determined for every plate with the time duration (Δt) that provides the highest Z’-factor. For each compound, when the MSD_Δt_ ratio was beyond the average + 3 SD of the MSD_Δt_ ratio for the negative control, the compound was regarded as causing a significant MSD change of EGFR and identified as a hit compound. Fig. 2c shows the selected 53 compounds with MSD_Δt_ ratios for compounds normalized to the negative control ratio. One of the marketed EGFR-targeted drugs, genistein, which is a flavonoid and known to inhibit tyrosine kinases including EGFR, was excluded due to its IC_50_ (50% inhibitory concentration) being above 10 μM, which was beyond our experimental condition.

For a more detailed assessment of the selected 53 compounds, we measured the dose-dependent effects of the compounds on EGFR mobility with and without EGF (Supplementary Fig. 4). Compounds with significantly different MSD values at the minimum and maximum concentrations were selected. When half of the difference between the MSD (averaged over ∼30 cells) at the two concentrations exceeded the sum of their SD, the compound was regarded as a hit compound. As a result, 18 compounds were obtained, but one of them, auranofin, was excluded from further analysis due to its cytotoxicity (cell death) during the treatment. Although these compounds showed similarly high MSD ratios, we found two types of effects on the MSD: one changed the receptor mobility primarily after the EGF addition (Fig. 2d), and the other before the addition (Fig. 2e). Among the 17 compounds, 10 compounds corresponded to the former case and suppressed the EGF-dependent decrease in a dose-dependent manner (Fig. 2d). These 10 were all kinase inhibitors: afatinib, erlotinib, OSI-420, gefitinib, lapatinib, and lapatinib ditosylate are known as EGFR-TKIs^8,22^; ponatinib and vandetanib are pan-TKIs^23^; and dasatinib and ibrutinib have been reported as Bruton’s tyrosine kinase inhibitors^24^ and can interfere with EGFR phosphorylation. All these EGFR-TKIs in the library were hit compounds (Supplementary Table 3), proving that our method was valid for exploring drugs targeting tyrosine phosphorylation of the receptor.

The other 7 hit compounds did not affect the EGF-induced decrease in mobility of EGFR but did affect the mobility without EGF stimulation (Fig. 2e). The reported drug actions and approved applicable symptoms are not apparently concerned with EGFR tyrosine kinases: broxyquinoline is an anti-infective agent; daunorubicin is an anthracycline antibiotic used for chemotherapeutic agents against cancer by inhibiting DNA topoisomerase II^25^; eltrombopag is used to treat severe aplastic anemia and activates thrombopoietin receptor (TPOR) to facilitate platelet production^26^; glafenine is a non-steroidal anti-inflammatory drug (NSAID) that inhibits cyclooxygenase-2/prostaglandin E2 (COX-2/PGE2) signaling^27^; nilotinib and sorafenib are not EGFR-TKI but TKI that largely inhibit EGFR-independent pathways and used respectively to treat primary kidney cancer^28^ and chronic myeloid leukemia^29^; and, finally, curcumin interferes with tyrosine kinases including EGFR and is approved as a food additive. Because these compounds induced EGFR mobility changes without EGF, they are suggested to interact directly or indirectly with EGFR by some actions other than the inhibition of tyrosine kinase. The effects of these compounds on EGFR are described in more detail below.

### EGFR clustering-based screening

Single-molecule tracking can obtain the brightness of each EGFR-mEGFP fluorescent spot, which reflects the number of fluorescent GFP involved. Uniform illumination was required to quantify the brightness. AiSIS achieved uniform TIR illumination over 3,000 μm^2^. Because CHO-K1 cells express no endogenous EGFR, their brightness typically reflects the oligomer/clustering size. The EGF concentration correlated with the brightness (Fig. 3a and b), indicating that monomers/dimers were relatively reduced with increasing concentration and larger size clusters were gained. The formation of clusters with integrating monomers/dimers has been suggested to correlate with the activation of downstream signaling through interactions with adaptor molecules^15^. Thus, the increase in brightness could be used as a screening index to assess signal transduction.

**Fig.3.**
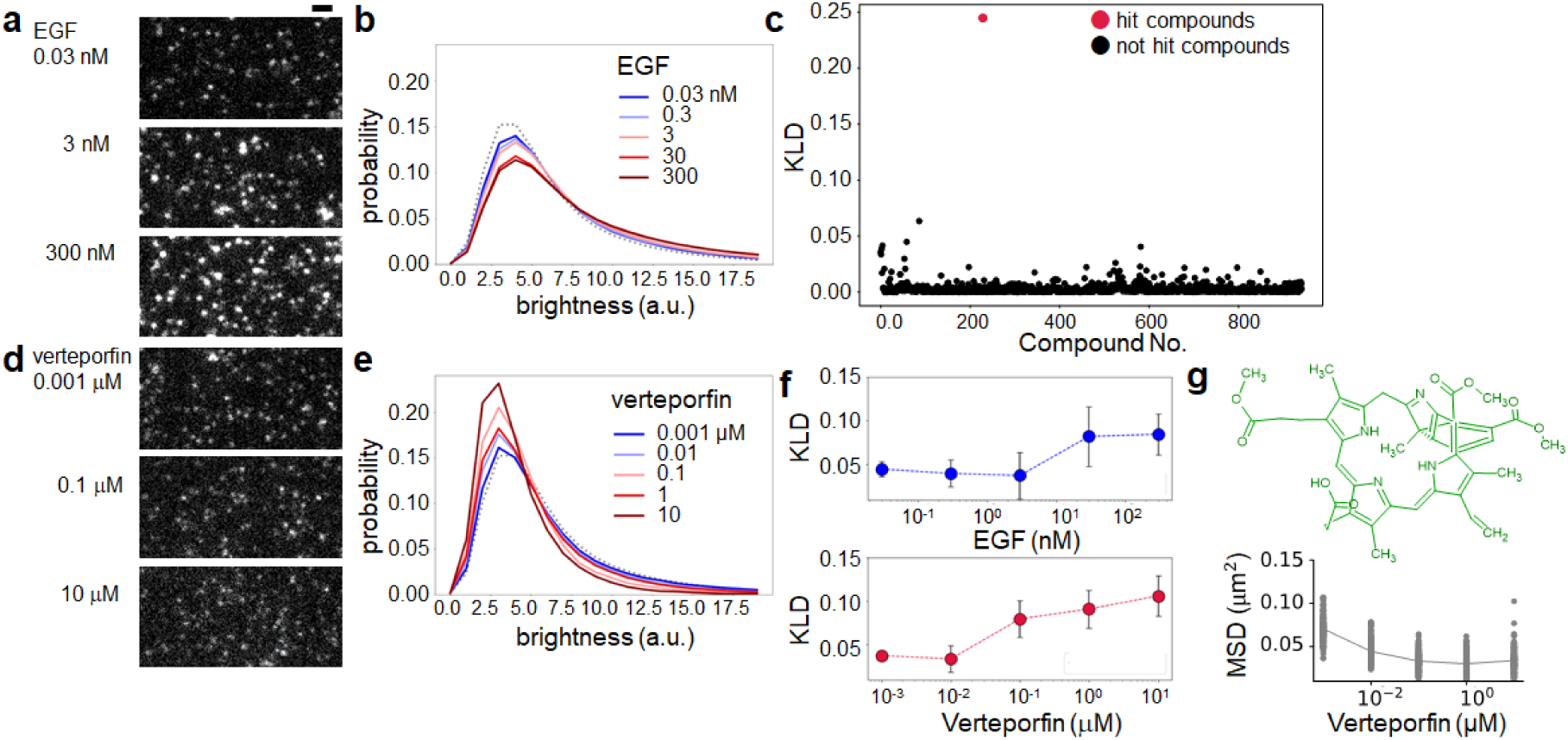
EGFR clustering-based screening for compounds in the FDA-approved library. **a,** EGF facilitated the formation of EGFR oligomers. Single-molecule images of EGFR showed brighter and larger fluorescent spots were increased as the EGF concentration increased. Single-molecule imaging began 1 minute after the EGF addition. Scale bar, 2 μm. **b,** The probability distribution of EGFR spot brightness. EGFR fractions shifted from small to large oligomers with higher EGF concentration. **c,** Effects of compounds on EGFR oligomerization were evaluated using the KLD of the brightness histograms before and after the compound treatment. Verteporfin exhibited extremely high KLD values (red dot). **d,** Verteporfin treatment increased the EGFR spots with lower brightness. **e,** The probability distribution of EGFR spot brightness. The fraction of smaller EGFR oligomers was increased with verteporfin concentration, suggesting disassembly of the EGFR oligomers. **f,** EGF (top) and verteporfin (bottom) dose-dependent KLD against the untreated condition. Error bars, SD. **g**, The dose-dependency was further evaluated by referring to MSD_167ms_ for verteporfin. Black circles denote MSD values without EGF, showing the reduction of MSD without EGF.

For the obtained single-molecule tracking data, we compared the probability density distributions of the spot brightness before and after compound treatment without EGF. The Kullback-Leibler divergence (KLD) was used to quantify the similarity of the histogram profiles. In this case, Z’-factors were calculated using KLD before and after the EGF addition (without any compound) as respective negative and positive controls. The observed Z’-factors did not satisfy the threshold of 0.5 due to the broad SD (Supplementary Fig. 3b). Nevertheless, the highest KLD value among the compounds (Fig. 3c) was obviously different from the KLD calculated for the negative control. This compound, verteporfin, diminished the number of brighter spots in a dose-dependent manner (Fig. 3d to 3f), suggesting that larger clusters of EGFR decreased on the plasma membrane. Verteporfin reduced EGFR diffusion in a dose-dependent manner without EGF, although it was not selected from the mobility-based screen due to its modest effects (Fig. 3g). Verteporfin is a photosensitizing drug for age-related macular degeneration^30^. As described in more detail below, the effect of this compound on EGFR was due to EGFR internalization and not to any photophysical effect.

### Characterization of hit compounds on EGFR dynamics, signal transduction and cell viability

To determine whether these non-EGFR-TKI compounds affect EGFR-dependent signaling in CHO-K1 cells expressing EGFR, which were used in the screening, we examined the activation and expression levels of both EGFR and the downstream signaling protein ERK (Fig. 4a-4c). Treatment with any of five non-EGFR-TKI compounds (broxyquinoline, daunorubicin, eltrombopag, sorafenib, and verteporfin) caused no significant inhibition of EGFR phosphorylation upon EGF stimulation in comparison with the control (compound-untreated) condition (Fig. 4a and 4b, left), a finding consistent with these compounds not being EGFR-TKIs. In the subsequent signaling, the phosphorylation of ERK fell upon treatment with any of the five compounds, especially eltrombopag, sorafenib, and verteporfin, although the expression levels of ERK were not affected, indicating an inhibition of EGFR-dependent signaling (Fig. 4a and 4b, right). The total amount of EGFR was decreased by treatment with the five compounds (Fig. 4c) regardless of EGF stimulation, indicating the destabilization of EGFR. Furthermore, EGFR was internalized in the cells treated with the five compounds without EGF, as shown in timelapse TIRF images (Fig. 4d) and by quantification (Fig. 4e). In the case of verteporfin, the photoreactive damage of GFP induced by repetitive laser irradiation was observed because of the photosensitizing effect of verteporfin (Supplementary Fig. 5a). Further observations of EGFR-mEGFP at the condition minimizing the photophysical effects suggested EGFR internalization (Fig. 4d, 4e, and Supplementary Fig. 5b, 5c). The examination by immunofluorescence microscopy of verteporfin supported EGFR internalization, as described later. In contrast with these five non-EGFR-TKI compounds, when treated with the other two compounds, glafenine and nilotinib, the cells exhibited no reduction of EGF-induced EGFR phosphorylation or downstream ERK (Fig. 4b) and showed no obvious internalization or degradation of EGFR (Fig. 4c and 4e). Thus, the five non-EGFR-TKI compounds (broxyquinoline, daunorubicin, eltrombopag, sorafenib, and verteporfin) caused the downregulation of EGFR-related signal transduction with internalization, even though EGFR phosphorylation was unchanged. That is, compounds that can alter EGFR signaling outside tyrosine phosphorylation were successfully selected by the single-molecule tracking-based screening method.

**Fig. 4.**
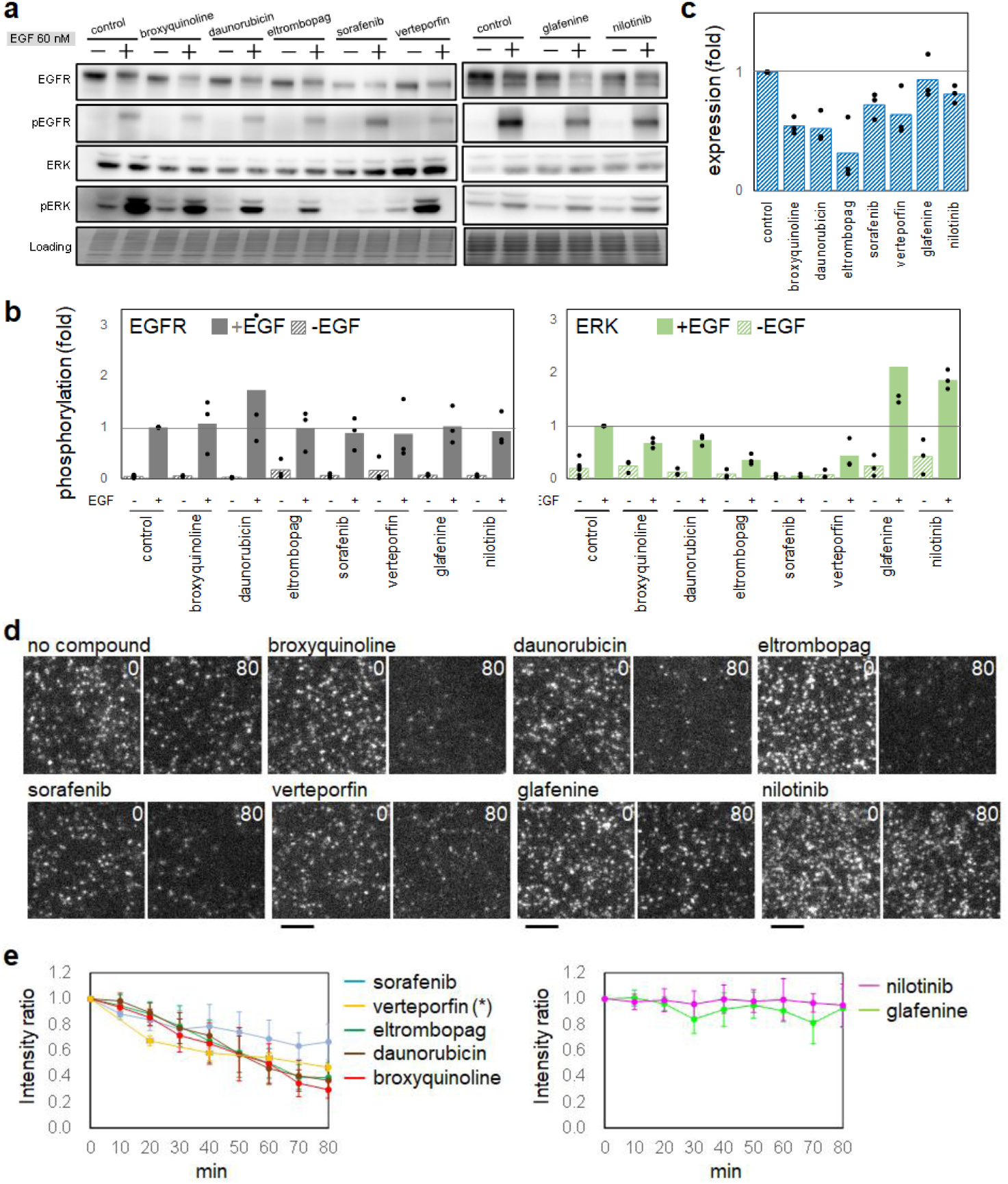
Effects of compounds on EGFR behavior and cell responses. **a-c,** Phosphorylation of EGFR and its downstream signaling protein (a and b) and EGFR protein expression in CHO-K1 cells expressing EGFR (c). After treatment with the indicated compounds for 1 hour (dashed bar), the cells were stimulated with EGF for 5 minutes (filled bar). Band densities were normalized to the EGF-stimulated (b) and unstimulated (c) condition. Gray lines indicate phosphorylation and expression levels in untreated cells. **d,** Timelapse single-molecule images acquired at 0 and 80 min after the compound treatment. Scale bar, 3 μm **e,** Quantified fluorescence per unit area, which was normalized to that at time 0. Solid lines correspond to the indicated compound. Timelapse images were acquired at the denoted time points in the same cells except verteporfin (*) for which the images were obtained at two time points (0 and the other) to avoid the fluorescence bleaching induced by multiple laser irradiation. Error bars, SD.

We next evaluated the effects of the hit compounds on EGFR-dependent cellular viability by utilizing various cell types that express EGFR because they are expected to depend on EGFR signaling for their survival: A431 and HeLa cells, which express endogenous EGFR, and EGFR-transfected Ba/F3 cells. We also utilized Ba/F3 and CHO-K1 cells, which do not express EGFR, as controls. Viability assays were carried out under 10 μM of compound for 72 hours. The left panel in Fig. 5a shows that all hit EGFR-TKIs (i.e. afatinib, erlotinib, gefitinib, lapatinib, ponatinib, vandetanib, dasatinib and ibrutinib) reduced the viability of A431 cells and EGFR-transfected Ba/F3 cells, consistent with a dependency on EGFR activity for their survival^31,32^. HeLa cells, which expressed EGFR 10-fold less than A431 cells^33^, also exhibited less viability with EGFR-TKI treatment, although they were somehow resistant to erlotinib and gefitinib, consistent with previous reports^34–36^. Similar to the effects of most EGFR-TKIs, the five non-EGFR-TKI compounds (broxyquinoline, daunorubicin, eltrombopag, sorafenib, and verteporfin) suppressed the viability of the three EGFR-expressing cell types, while the other two non-EGFR-TKIs (glafenine and nilotinib) caused higher viability. In the right panel in Fig. 5a, the viability of CHO-K1 and Ba/F3 cells was not significantly suppressed by EGFR-TKIs or the five non-EGFR-TKI compounds. These observations indicate that the five non-EGFR-TKI compounds suppress EGFR-dependent cell survival. The viability of CHO-K1 cells lacking EGFR expression was sensitive to ponatinib and dasatinib, possibly due to these compounds acting as TKIs for other tyrosine kinases as well as EGFR.

**Fig. 5.**
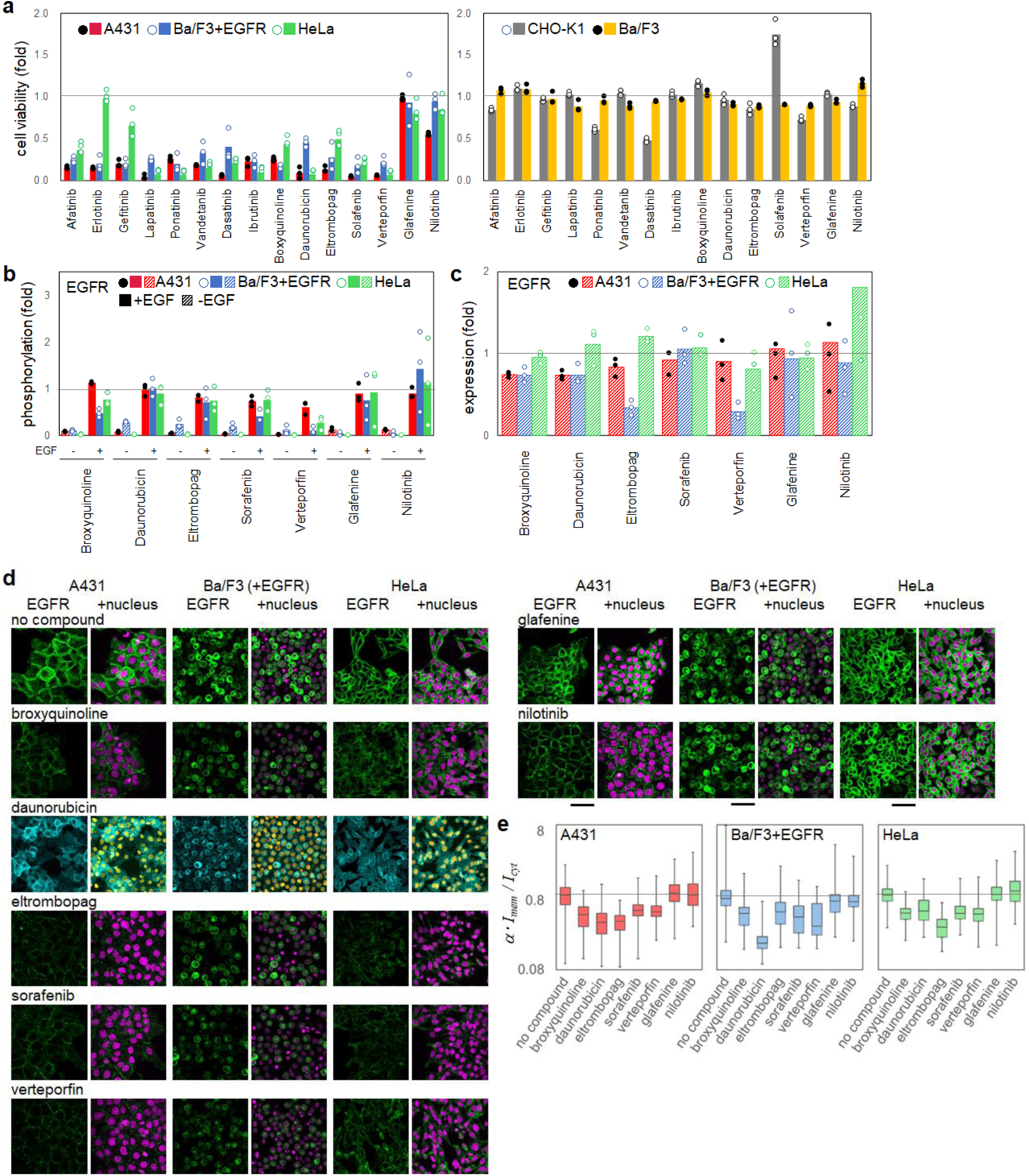
Effects of compounds on different cell lines. **a,** Viabilities of cell lines treated with the hit compounds including EGFR-TKIs. Left, EGFR-expressing cell lines: A431 (red), EGFR-transfected Ba/F3 (light blue), and HeLa (green). Right, non-EGFR-expressing cell lines: CHO-K1 (gray) and Ba/F3 (orange). Viabilities were normalized to those of compound-untreated control cells. **b** and **c,** Phosphorylation of EGFR (b), which was normalized by that in cells after EGF stimulation, and EGFR protein expression (c), which was normalized by that in cells before the stimulation. Compound treatment was carried out with the indicated compounds for 1 hour (dashed bar), and cells were subsequently stimulated with EGF for 2 minutes (filled bar). Gray lines indicate phosphorylation and expression levels in compound-untreated cells. **d,** Immuno-fluorescent images of EGFR in A431, Ba/F3 expressing EGFR, and HeLa cells acquired at 1 hour after the compound treatment. Each cell was recognized by its nucleus. Because daunorubicin has an auto-fluorescence in the nucleus at 488 nm excitation, EGFR was observed using a second antibody at 640 nm excitation. For the other compounds, EGFR was observed at 488 nm excitation (see Methods). Scale bar, 100 μm **e,** Internalization was quantified using the average fluorescence intensities of the plasma membrane and cytoplasm regions (α•*I*_mem_ */ I*_cyt_, see Methods). Values were normalized to that of untreated cells (gray lines). The box- and-whisker plots show the median as horizontal lines, the first and third quartiles as box ends, and the whiskers as minimum and maximum values.

We further evaluated the effects of the five non-EGFR-TKI compounds on EGFR signaling in the various cell lines. The EGF-induced phosphorylation of EGFR was observed with the non-EGFR-TKI compounds in A431, HeLa, and EGFR-transfected Ba/F3 cells (Fig. 5b and Supplementary Fig. 6), although at varying degrees especially for verteporfin. The expression level of EGFR tended to be reduced slightly by the five compounds in A431 and EGFR-transfected Ba/F3 cells but rarely in HeLa cells (Fig. 5c). Verteporfin, in particular, caused EGFR-destabilization in EGFR-transfected Ba/F3 cells, which can explain the reduced EGF-induced EGFR phosphorylation (Fig. 5b).

The stability and internalization of EGFR were further examined in the EGFR-expressing cell lines by immunofluorescence microscopy, in which the photophysical effect of verteporfin need not be considered. Treatment with any of the five non-EGFR-TKI compounds caused a decrease in EGFR localization on the membrane in A431, HeLa and EGFR-transfected Ba/F3 cells compared to untreated cell lines (Fig. 5d). In particular, eltrombopag, sorafenib, and verteporfin tended to decrease the total expression level of EGFR, suggesting the internalization and degradation of EGFR. Fig. 5e shows the ratio of EGFR localization on the plasma membrane and in the cytoplasm of the EGFR-positive cell lines treated with the non-EGFR-TKI compounds, revealing the five compounds enhanced the internalization of EGFR. In contrast, the other two compounds (glafenine and nilotinib) did not alter EGFR-dependent cell viability (Fig. 5a), EGF-induced EGFR phosphorylation (Fig. 5b), the expression level of EGFR (Fig. 5c), or the membrane localization of EGFR (Fig. 5d and 5e), indicating no effects on EGFR-dependent signaling or cell survival, although these compounds affected EGFR mobility (Fig. 2e). Thus, the hit compounds, which were selected using EGFR-expressing CHO-K1 cells, were confirmed to robustly exert inhibitory effects on the EGFR-dependent cell viability by inducing the internalization and degradation of EGFR in various cell types including A431, EGFR-transfected Ba/F3, and Hela cells, albeit to varying degrees.

Overall, the hit EGFR-TKIs that inhibited the phosphorylation-dependent mobility shift of EGFR reduced the EGFR-dependent viability. The five non-EGFR-TKI compounds (broxyquinoline, daunorubicin, eltrombopag, sorafenib, and verteporfin) that decreased EGFR diffusion or clusters without EGF, reduced EGFR-dependent cell viability by downregulating EGFR. In contrast, the other two non-EGFR-TKI compounds (glafenine and nilotinib) that affected EGFR mobility without EGF stimulation did not affect the EGFR-dependent signaling needed for cell survival.

## Discussion

Here, we demonstrated single-molecule tracking-based drug screening to explore compounds effective against EGFR. In addition to the methods used in previous single-molecule screening for nuclear receptors^37^, AiSIS enabled highly precise auto-focusing on the plasma membrane receptor and uniform TIR illumination for the quantification of fluorescence brightness. Our method observed changes in EGFR mobility and clustering instead of focusing on a specific process of the signaling, allowing it to detect changes in multiple processes dependent on the ligand. Because changes in the mobility and cluster distribution of the target molecule can hardly be measured by biochemical methods but can be observed by single-molecule imaging as signaling processes, our single-molecule tracking-based screening has the potential to detect compounds that cannot be detected by conventional methods. Screening by assessing several treatment conditions of ligand/compound led to a wide selection of compounds affecting EGFR functions that are related or unrelated to EGF stimulation (Fig. 2). Among the hit compounds, those suppressing the EGF-induced mobility shift of EGFR were identified as EGFR-TKIs, proving the reliability of the method for drug screening. One of the possible reasons why EGFR-targeted TKIs can be detected by mobility-based screening is described in Fig. 6a. Accumulating evidence shows that EGFR undergoes structural changes during its phosphorylation upon EGF stimulation and then translocates to a confined membrane region via interactions with membrane components (lipid, proteins, etc.), leading to slower receptor mobility^15^. Changes in the MSD of EGFR were proven to couple tightly with the phosphorylation in a dose-dependent manner (Fig. 1f). The competitive docking of TKI to the ATP-binding pocket of EGFR prevents the receptor from subsequent structural changes during phosphorylation, which thus suppresses the decrease in mobility and subsequent signaling processes.

**Fig. 6.**
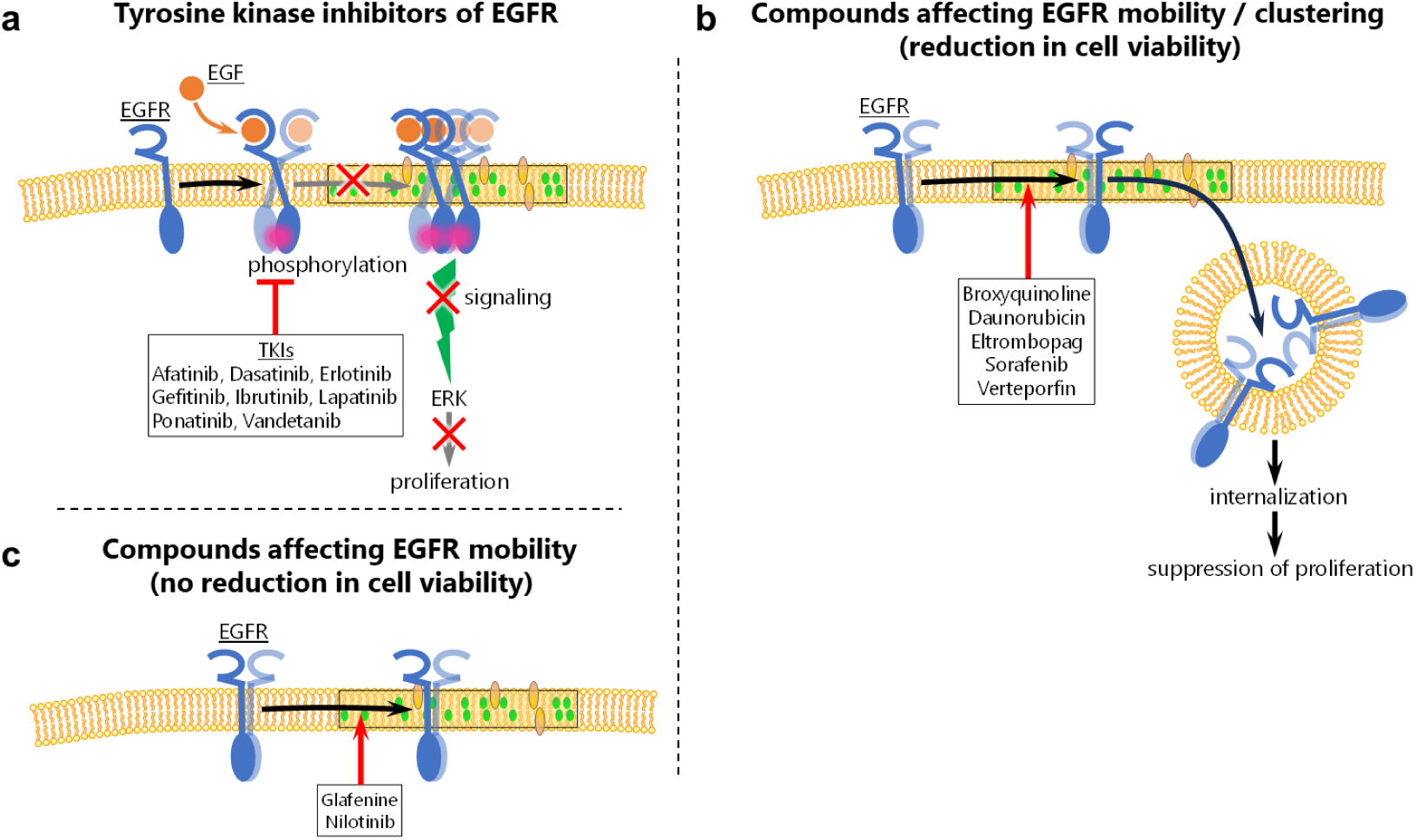
Categorized schemes for effects of hit compounds on EGFR. Observed schemes of the hit compounds. **a,** TKIs suppress the EGF-induced phosphorylation of EGFR, thereby inhibiting the subsequent signaling process. **b,** Compounds reducing both EGFR mobility and clustering induce EGFR internalization. **c,** The internalization of EGFR was not observed in cells treated with compounds that reduced EGFR mobility but not cell viability (glafenine and nilotinib).

In addition to the detection of EGFR-TKIs, our screening detected non-EGFR-TKI compounds (i.e. broxyquinoline, daunorubicin, eltrombopag, sorafenib, verteporfin, glafenine, and nilotinib), none of which are known to act on EGFR directly. Fig. 6b illustrates a scheme for the broxyquinoline, daunorubicin, eltrombopag, sorafenib, and verteporfin effects; these compounds commonly reduced mobility, induced internalization, decreased the expression levels of EGFR, and affected EGFR-dependent signaling and viability. We assume that these compounds force EGFR to a specific membrane subdomain where receptor mobility is lower. EGFR in the membrane subdomains may internalize via caveolin-mediated endocytosis^38,39^. The internalization exerts a harmful effect on cell viability by suppressing EGFR signaling for proliferation and survival^40^. For example, oxidative processes in cells are induced by various compounds (e.g. broxyquinoline) and have been suggested to affect EGFR signaling^41^. Previous studies have shown that broxyquinoline stabilizes hypoxia inducible factor (HIF)-1^42^ to accumulate caveolin-1 in the lipid raft to constitute caveolae^43^, which directly interacts with EGFR, suggesting the confined mobility and internalization of the protein complex including EGFR. In fact, we confirmed that broxyquinoline increased HIF-1 and the internalization of EGFR with caveolin in EGFR-transfected CHO-K1 cells (Supplementary Fig 7). Furthermore, some compounds may induce complex formation including EGFR, thereby promoting the internalization and degradation of EGFR^38,44,45^. In addition, some compounds are reported to interact with the plasma membrane to possibly alter EGFR mobility^46,47^. Fig. 6c illustrates the case of glafenine and nilotinib. No significant change was observed in the EGFR-dependent viability of cells treated by these two compounds (Fig. 5). Additionally, the phosphorylation of downstream proteins was not suppressed, and no internalization/reduction of EGFR expression occurred (Fig. 4 and 5), indicating no downregulation of EGFR signaling by these two compounds.

As described above, the hit compounds obtained in the present study were categorized into three types based on the observed EGFR behavior and cell responses, with each type suggested to depend on a different mechanism of action. The scheme in Fig. 6a, which illustrates that EGFR-TKIs can be obtained by the mobility-based screening, is useful for searching for novel TKIs if EGFR mutants resistant to existing TKIs appear to change their motility upon EGF-stimulated activation^48^. The scheme in Fig. 6b, which was suggested from the five non-EGFR-TKI compounds, is effective for removing pathogenic cells that excessively express EGFR, a common phenotype of various cancers, by inducing cell death following EGFR internalization. Therefore, drug repositioning for novel EGFR drugs may be possible using compounds that exhibit this scheme. In Fig. 6c, compounds that do not significantly affect cell viability but exert an effect on receptor mobility might be drug seeds with little cytotoxicity. Besides EGFR, various types of receptors on the plasma membrane have been reported to change their mobility according to their activation and the cluster formation to propagate downstream signaling^15,37,49^, suggesting that single-molecule tracking-based drug screening is applicable to a wide variety of membrane receptors.

## Data availability

Source data are provided with this paper. All other data supporting the findings of this study are available from the corresponding author upon reasonable request.

## Code availability

All Python and ImageJ scripts supporting the findings of this paper are available upon reasonable request.

## Acknowledgments

We thank A. Kanayama for providing experimental support, P. Karagiannis for editing the manuscript, and all members of the Ueda lab for discussions. The epithelial-like cell lines, CHO-K1 (RCB0285), HeLa (RCB0007), A431 (RCB0202), and the mouse pro-B cell line, Ba/F3 (RCB4476), were provided by the RIKEN BRC through the National Bio-Resource Project of the MEXT, Japan. This project was supported by the Translational Science and Medicine Training Program, AMED, Center for Supporting Drug Discovery, Life Science Research, Osaka University, and the RIKEN Program for Drug Discovery and Medical Technology Platforms (DMP).

## Funding

AMED Grant Number JP23ym0126815, Japan; JST CREST Grant Number JPMJCR21E1, Japan; MEXT Grants-in-Aid for Scientific Research(B) (18H01839), Japan; MEXT Grants-in-Aid for Scientific Research(B) (22H02593), Japan; JSPS Grant-in-Aid for Scientific Research on Innovative Areas (18H05414), Japan.

## Author contributions

Conceptualization: MH and MU. Methodology: DW, MY, and MH. Investigation: DW and MH. Funding acquisition: MH and MU. Supervision: MU. Writing – original draft: DW, MH, and MU. Writing – review & editing: MH and MU.

## Competing interests

The authors declare no competing interests.

## Data and materials availability

Data availability: All data are available in the main text or the supplementary materials. All code used in the analysis are available from the authors upon reasonable request.

## Notes

### Competing Interest Statement

The authors have declared no competing interest.

### Summary of Updates

Revised text, figures, citations, and supplementary materials to clarify. No change to manuscript substance.

