## Supplementary Materials for "Single-molecule tracking-based drug screening"

#### Methods

##### Cell preparation

CHO-K1, A431, and HeLa cells (RIKEN BRC through the National Bio-Resource Project of the MEXT, Japan) were grown at 37°C in Ham’s F-12 medium (CHO-K1) or Dulbecco's modified Eagle's medium (A431, HeLa; DMEM, FUJIFILM Wako Pure Chemical, Japan); both media were supplemented with 10% fetal bovine serum (FBS). Ba/F3 cells (RIKEN BRC) and Ba/F3 cells expressing EGFR[50,51] (a kind gift from Dr. Ryo Iwamoto and Dr. Eisuke Mekata, Osaka University) were cultured at 37°C in RPMI medium supplemented with 10% FBS and 4 ng/mL mouse IL3 (091-03971, Fuji-Wako, Japan). In the case of Ba/F3 cells expressing EGFR, 20 ng/mL EGF (315-09, PeproTech, USA) was added to the medium. The culture medium was replaced with DMEM minus phenol red or FBS for starvation two hours before single-molecule imaging, when the medium was changed to DMEM containing 5 mM PIPES. The cells were cultured until 90% confluence and observed in 60 wells of a 96-well plate (GP96000, Matsunami Glass, Japan); the peripheral wells were excluded to avoid interference between the microscope stage and the objective lens.

##### Automated in-cell single-molecule imaging system (AiSIS)

AiSIS consists of an automated microscope for single-molecule imaging and robotics for liquid dispensing. Total internal reflection (TIR) illumination was configurable with a high-magnification objective, PlanApo 60X NA 1.49 (Nikon, Japan), to acquire single-molecule images at the basal cell surface under an inverted microscope (Ti2-E, Nikon, Japan). Auto Imaging System (ZIDO corp., Japan), including an autofocus device and AI-aided cell searching applications, was equipped to automatically obtain in-focus images of the cells with a suitable density of fluorescent spots for single-molecule analysis. Lasers at wavelengths of 488 nm (OBIS 488LS, Coherent, USA) were used to excite GFP fused with EGFR. The dichroic mirror/emission filter set was DM495/BA500-545 (Nikon, Japan). The images were acquired at a frame rate of 33 ms for 25 frames using a sCMOS camera (ORCA-Flash4.0 V2, Hamamatsu, Japan). A liquid-handling robot (Cavro Omni Robot, Tecan, USA) automatically sucked 100 μL of two-fold the final concentration of EGF solution from the well in the storage plate and dispensed the solution into the target well of the cell culture plate with 100 μL of observation medium.

**Autofocus device**

The device consists of a light source, magnifier optics, a slit located at the plane optically conjugate to the basal cell surface, a CCD camera for acquiring the slit image, and a control unit that feedback-controls the position of the objective lens (Fig. 1b). An 830-nm laser is used as the light source, and its incident angle is switched between two values using a Galvano mirror with a frequency of 10 Hz. The slit image on the CCD is moved according to the position of the objective lens. When the objective is located at an under- or over-focus position, the image is seen at the opposite side across the center, where it is formed by the objective at the in-focus position (Supplementary Fig. 1a). The center can be identified quickly as the middle point between two different slit images obtained by switching the incident angle of the laser 180 degrees. The side of the image corresponds to the z-direction of the shift of the objective from the in-focus position. The calculated deviation from the center is transmitted to a control unit to feedback-control the objective position. The gap difference between the in-focus positions determined by the device and human eyes can be adjusted by setting an offset value. Supplementary Fig. 1b shows the flow of the autofocusing process. This method, using two images with light from exactly opposite directions, is robust for variations in the sample condition (e.g., refractive index) that affect the optical path at the sample-coverslip interface and cause inaccurate focusing by conventional methods. The device can also be applied to normal microscopy, especially with high magnification. In Supplementary Fig. 1c, another method for switching the laser direction is introduced. A circular transparent plate with a thin circular coating (e.g., aluminum) forms thin light-nontransparent regions inside and outside each half of the circle. The laser illumination goes through the circles; therefore, when the plate rotates, the optical path is switched according to the rotational frequency, enabling illumination from opposite directions.

**Single-molecule tracking**

Single-molecule tracking was carried out on the acquired images to obtain the positions and brightness of fluorescent EGFR-mEGFP using commercial software (Auto Analysis Software, AAS, ZIDO, Japan). The first process of the tracking was to determine bright spots in the image by selecting square regions with 11 x 11 pixels beyond a threshold for cross-correlation between pixel values and the two-dimensional Gaussian distribution. Then, the spot region was fitted with the following function:

$$I(x, y; I_0, x_g, y_g, \sigma_A, a, b, I_{back}) = I_0\, exp\left[-\frac{(x-x_g)^2+(y-y_g)^2}{2\sigma_A^2}\right] + a(x - x_g) + b(y - y_g) + I_{back},$$

where $x$ and $y$ denote the pixel positions. The distribution of the pixel values was expressed as a Gaussian function with a peak intensity of $I_0$ at the centroid, $(x_g, y_g)$ and a variance of $\sigma_A{}^2$, plus a background with an inclination described by $a$ and $b$ above the offset intensity $I_{back}$. A single-molecule trajectory was generated by connecting the spots as follows: all possible connections between two spots at times $t$ and $t - 1$ with a center-to-center distance below 6 pixels were listed, and the shortest connection was selected. Trajectories outside the cell region determined by the AI-aided cell search algorithm were removed.

**Molecular mobility analysis**

The mobility of lateral diffusion was analyzed with the MSD calculated from the positions of the fluorescent spots with the following equation:

$$MSD(n\Delta t) = [\{x_i(n\Delta t + m\Delta t) - x_i(m\Delta t)\}^2 + \{y_i(n\Delta t + m\Delta t) - y_i(m\Delta t)\}^2]_{i,m} \cdot$$

Here, $x_i$ and $y_i$ represent the single-molecule position in the i-th track, $n$ and $m$ denote the frame number, $\Delta t$ is the time interval between frames (33 ms), and $[\ ]_{i,m}$ denotes the average over $i$ tracks and $m$ frames. For Fig. 1f and 1g, MSD at $\Delta t = 500$ ms was used. The dose-response curve (Fig. 1f) was plotted as the MSD ratio and fitted using the following equation to calculate the $EC_{50}$:

$$MSD = MSD_{max} - \frac{MSD_{max} - MSD_{min}}{1 + (\frac{EC_{50}}{[L]})^h},$$

where $MSD_{max}$ and $MSD_{min}$ are the MSD ratios of the upper and lower boundaries, respectively, $[L]$ is the ligand (EGF) concentration, and $h$ is the Hill coefficient. To display the dose-response curve in comparison with the phosphorylation level, the MSD ratio was converted as follows:

$$MSD_{converted} = \frac{MSD_{max} - MSD}{MSD_{max} - MSD_{min}}.$$

**Compounds**

The library of FDA-approved compounds was provided in 96-well plates from the Center for Supporting Drug Discovery and Life Science Research, Osaka University. 3 μL of 1 mM compound in DMSO was dispensed in each well and diluted to 10 μM with DMEM before screening. The medium in 60 wells of the 96-well plate with cultured cells was replaced to the compound solution and incubated at 37°C for 1 hour. Both sides of the compound-treated wells (total 2 x 6 wells) were used for the positive and negative controls (with and without 10 μM gefitinib), respectively.

**Single-molecule screening**

For the compound screening, single-molecule imaging was executed on 20 cells before and 20 cells after EGF stimulation. 100 µL of 120 nM EGF was added to every well (final concentration, 60 nM). The MSD ratio was used to confirm the quality of the screening and select the hit compounds. For each plate, we calculated the Z'-factor using the MSD ratios of the wells with the positive (10 µM gefitinib in DMSO) and negative (DMSO only) controls. The MSD at Δt that provided the best Z'-factor in each well plate was used for Fig. 2c and at 167 ms for Fig. 2d and 2e. The average and SD of the MSD ratio of positive and negative controls were used to calculate the Z'-factor.

$$Z' = 1 - \frac{(3 \times SD_{positive}) + (3 \times SD_{negative})}{Avg_{positive} - Avg_{negative}}.$$

Here, *Avg* and *SD* respectively represent the averages and SD of the MSD for the positive and negative controls. In Fig. 2c, the MSD ratio for each compound was normalized to that of the negative control, and any compound with a ratio greater than the sum of the average and three-fold SD of the negative control was defined as a hit compound. For the compound-dose dependent assay shown in Fig. 2d and 2e, cells were treated with 100 µL of 10 µM, 1 µM, 0.1 µM, 0.01 µM, and 0.001 µM compound solution for 1 hour before single-molecule imaging. Regardless of the EGF addition, when the MSD value was larger than half the difference of the MSD at the minimum and maximum compound concentration plus their SD, the compound was recognized as effective for EGFR.

**Cell viability assay**

A431 cells in 96-well plates (1860-096, Iwaki, Japan) were incubated in 10 µM compound solution at 37℃ for 72 hours. Then, the medium was replaced with 10 µL of Cell Counting Kit - 8 (Dojindo, Japan) diluted in 100 µL HBSS. After 2 hours of incubation, absorption of the medium was measured for each well with a wavelength of 450 nm using a plate reader (Infinite F50 Plus, Tecan, USA).

**Western blotting**

After the compound and/or EGF treatment, the cells were lyzed in SDS sample buffer. The lysate was electrophoresed on a 10% SDS-polyacrylamide precast gel (192-14961, SuperSep Ace, 10%, 17well, FUJIFILM Wako Pure Chemical, Japan) Then, the proteins were transferred to a 0.45 μm pore PVDF membrane (034-25663, ClearTrans PVDF Membrane, Hydrophobic, 0.45μm, FUJIFILM Wako Pure Chemical, Japan) and reacted with antibodies against the following targets: EGFR (#4267, Cell signaling technology (CST), USA), phospho-EGFR (Tyr1068, #4267, CST, USA), AKT (#9272, CST, USA), phospho-AKT (Ser473, #4060, CST, USA), ERK1/2 (#9107, CST, USA), and phospho-ERK1/2 (Thr202/Try204, #9106, CST, USA). For chemiluminescent antibody detection, HRP-linked anti-rabbit IgG (#7074, CST, USA) and HRP-linked anti-mouse IgG (#7076, CST, USA) were used with ECL prime reagent (Cytiva, USA). The loading control containing proteins ranging from 37 to 250 kDa was obtained from the SDS gel stained with CBB. Quantification of the band densities were carried out using ImageJ software (NIH). Two squares were set such that one square was placed in the band and another square was far away from the band, and the difference in the average intensities of these regions was defined as the band intensity. The intensity of the anti-phosphorylated antibody was normalized to that of the anti-protein antibody. The phosphorylation level was calculated by dividing the normalized intensity of the phosphorylated protein for each condition by that for 30 nM EGF without compound. The EGF dose-response curve was fitted with the Hill equation as follows:

$$phosphorylation = min - \frac{max - min}{1 + \left(\frac{EC_{50}}{[L]}\right)^{h}}.$$

Here, $h$ indicates the Hill coefficient, *max* and *min* denote upper and lower bounds, respectively, and $[L]$ is the concentration of the ligand, EGF. The protein expression amount was calculated by normalizing the intensity of the protein for each condition by that of the loading control.

**Internalization assay**

For quantification of the internalization (Fig. 4d and 4e), CHO-K1 cells expressing EGFR-mEGFP in a 96-well plate were treated in compound solution for 1 hour. The compound solutions were the same as those used in the experiment above. After the compound treatment, timelapse single-molecule images (Fig. 4d) were acquired by AiSIS at the same positions in a well every 10 minutes. For verteporfin, which showed rapid photobleaching of EGFR-mEGFP as a photophysical effect induced by repetitive laser irradiation, the images were obtained at the same positions at only two time points (0 min and 20, 40, 60 or 80 min) to minimize the effect. In the obtained images, the average brightness was measured for circled regions of interest (ROI) within a cell and outside the cell (background). The difference in the brightness between these regions was calculated for each time point and normalized to the value at time 0 (Fig. 4e). To consider the effect of fluorescence bleaching caused by repetitive irradiation by the laser, timelapse images of the cells without compound treatment were acquired and analyzed by the same method.

**Immunofluorescence microscopy**

The compound treatment of A431 and HeLa cells cultured in 24-well plates was carried out in 10 μM solutions for 1 hour at 37°C after the cells were incubated in DMEM without FBS for 6 hours. Subsequently, the compounds were washed out with HBSS, and the cells were fixed with 4% PFA for 30 min at 4°C and solubilized with 0.5% Triton X100 in HBSS at room temperature, followed by washing with HBSS three times and left at 4ºC overnight. After the blocking process with HBSS containing 2% BSA for 15 min, 1:300-diluted anti-EGFR antibody (#4267, CST,

USA) or 1:10000-diluted anti-caveolin antibody (#3267T, CST, USA) was added as the primary antibody for 1 hour at room temperature. The cells were washed with HBSS three times and labeled with 1:1000 anti-IgG antibody conjugated with Alexa 488 or Alexa 647 (#A-11034 or #A-21244, respectively, Invitrogen, USA) for 1 hour at room temperature. The nucleus was fluorescently labeled with 0.05% NucSpot Live 650 (Biotium, USA) for 30 min at room temperature. Observation of the samples was done by confocal microscopy (Nikon A1) with a 20X objective lens (Nikon PalnApo) operated with NIS elements software (Nikon). The preparation process for EGFR-transfected Ba/F3 cells was almost the same as that for other cells, except that floating cells in Eppendorf tubes were centrifuged with 800 g for 2 min to replace the supernatant after every step. For the observation, the suspension of cells in HBSS was in a 96-well plate. The obtained images were analyzed by quantifying the fluorescence intensity of the plasma membrane and cytoplasm to calculate the ratio of EGFR in these regions. The regions of the plasma membrane and cytoplasm were identified as the area within 5 pixels inside the cell boundary, which was determined with Cellpose 3.0, and the other pixels within the cell, respectively. The average fluorescence intensities of the plasma membrane ($F_{mem}$), the cytoplasm ($F_{cyt}$), and background area ($F_{bck}$) where no cells existed were obtained to calculate the ratio, $I_{mem} / I_{cyt} = (F_{mem} - F_{bck}) / (F_{cyt} - F_{bck})$, as the extent of internalization. In addition, the decrease in total EGFR in a cell due to degradation was taken into account as the intensity ratio of the whole cell with to without compound treatment $\alpha = (F_{drug} - F_{bck}) / (F_{ctrl} - F_{bck})$. The product of this ratio ($\alpha \cdot I_{mem} / I_{cyt}$) includes both the internalization and degradation effects of a compound, which correspond to the fluorescence change in the obtained image. The calculations were performed using a custom-made program written in Python 3.8.18.

#### Supplementary Figures

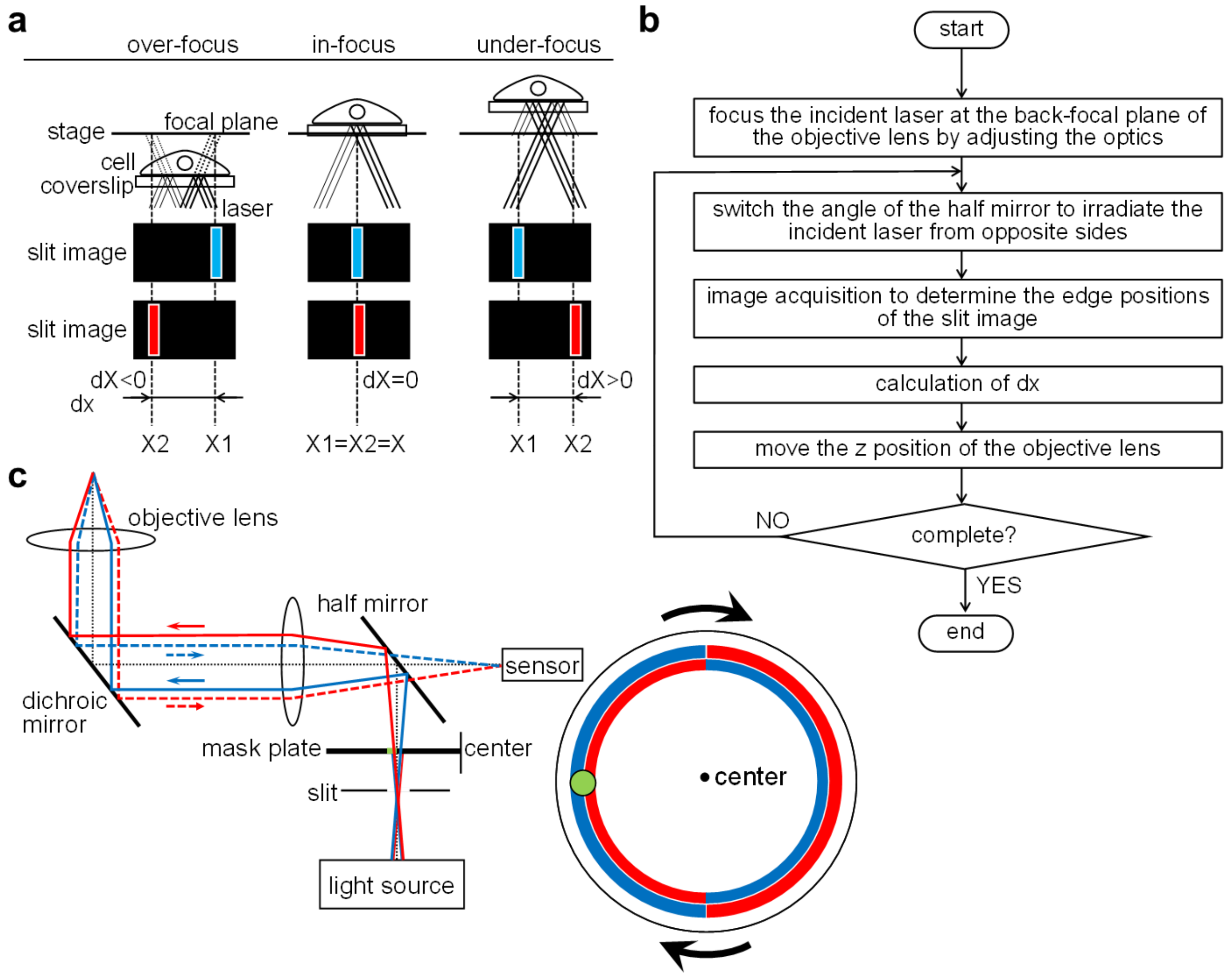


**Supplementary Fig. 1. Details of the autofocus device in AiSIS.**

**a,** Correspondence between the slit images generated with laser light in opposite directions and the focal positions. From under- to over-focus, these slit images (red and blue bars) exchange their locations through the in-focus position where the images are overlapped. **b,** A flowchart of the autofocus algorithm. **c,** Instead of switching the illumination, rotation of a mask-plate (right), in which light is transmitted at the green region but not at the blue or red regions, can be used. The blue region in the mask-plate indicates the red region rotated 180˚. The red and blue optical paths correspond to light affected by the same-colored regions in the mask-plate. Solid and dotted lines in the paths denote incident and reflected light, respectively.

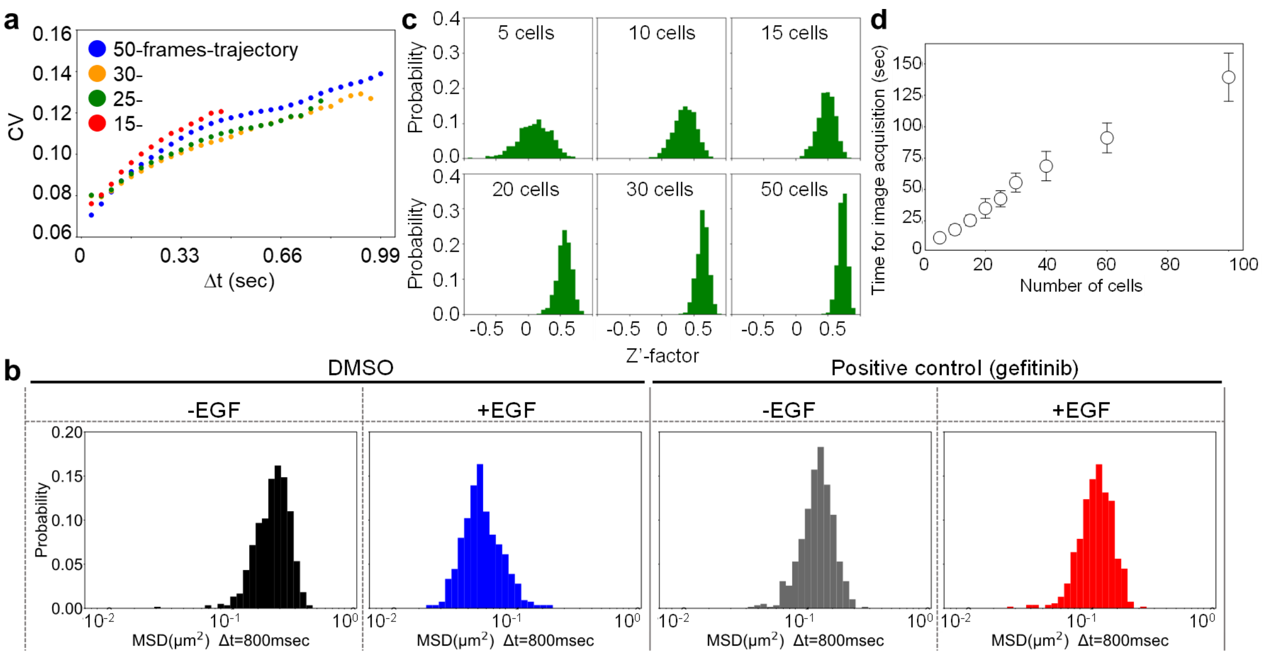


**Supplementary Fig. 2. Optimized conditions for image acquisition.**

**a,** The required number of frames in single-molecule imaging to obtain results with low deviation and high reproducibility. Single-molecule images were acquired for 50 frames at 30 frames/sec, and the first 15, 25, 30, and 50 frames were extracted from the same images to obtain the trajectories of single molecules. The MSD ratio at Δt before and after EGF addition was calculated for each trajectory group and obtained for each well where images from 10 cells both before and after EGF addition were acquired. The coefficients of variation (CV) of the MSD ratios were calculated from 48 wells. The MSD ratio from the trajectories of 15 frames showed a larger CV than the ratio from other imaging conditions due to the smaller number of frames, but the trajectories of 50 frames exhibited more GFP fluorescence bleaching. Therefore, movies of 25-30 frames (0.8 - 1.0 sec) were determined as suitable for the analysis. **b,** The MSD distribution at Δt = 800 ms obtained from 600 cells with and without EGF stimulation under gefitinib or DMSO (control) treatment. **c,** The required number of cells for single-molecule imaging to obtain a high Z'-factor. The Z'-factor at the indicted cell number was calculated by using MSD data selected randomly from the cell populations shown in b, and the statistical distribution of the Z'-factor was obtained. MSD ratios with and without gefitinib treatment, respectively, were used as positive and negative controls. For each graph, the calculation was done for 1000 different combinations of cells. **d,** The relationship between the number of measured cells and the elapsed time. Based on the assessment, 20 cells were measured before and after EGF addition in each well to obtain a Z'-factor satisfying the required screening accuracy (≥ 0.50) in as minimum time as possible.

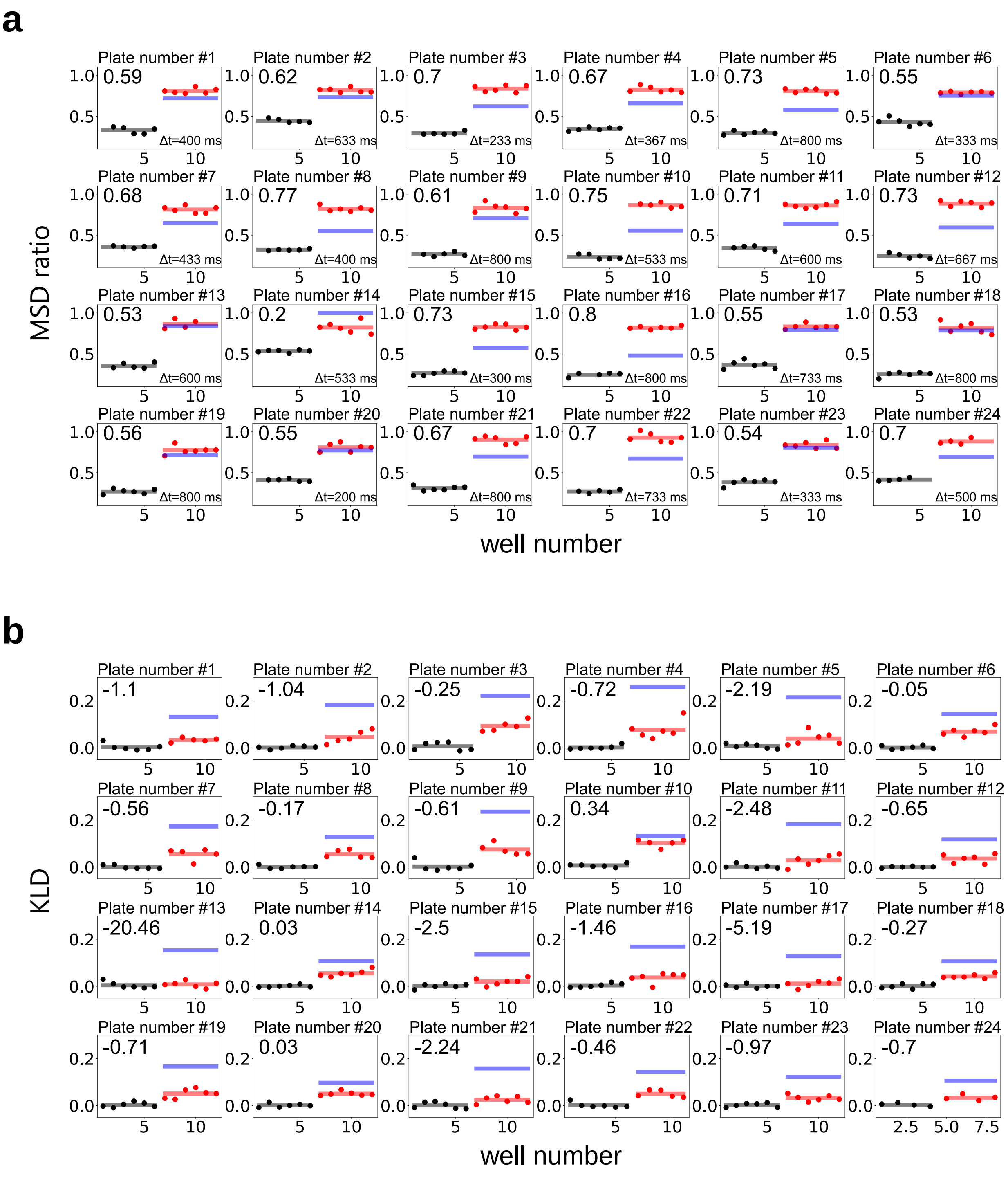


**Supplementary Fig. 3. Z'-factor values of each well plate for EGFR single-molecule screening.**

Z'-factor values of every well plate (upper left) obtained for mobility-based and clustering-based screenings, which were calculated from the MSD ratio and KLD, respectively. **a,** The MSD ratio is plotted without (black) and with (red) gefitinib treatment, which represent the negative control (cells treated only with EGF) and positive control (cells treated with gefitinib and EGF), respectively. Δt (bottom right) indicates the duration for the MSD calculation that provided the best Z'-factor for each well plate used in the screening. **b,** KLD before (black) and after (red) EGF addition. Each point was calculated from images of 10 cells in the same well. Red and black lines indicate the average of the plots. Blue lines show the required average of the positive control to provide a Z'-factor ($\geq 0.50$) sufficient for screening, which was calculated using the deviation of the positive/negative controls and the average of the negative control. Only well-plates with red lines above the blue line were used for all subsequent analysis. For example, plate number #14 was excluded from the mobility-based screening.

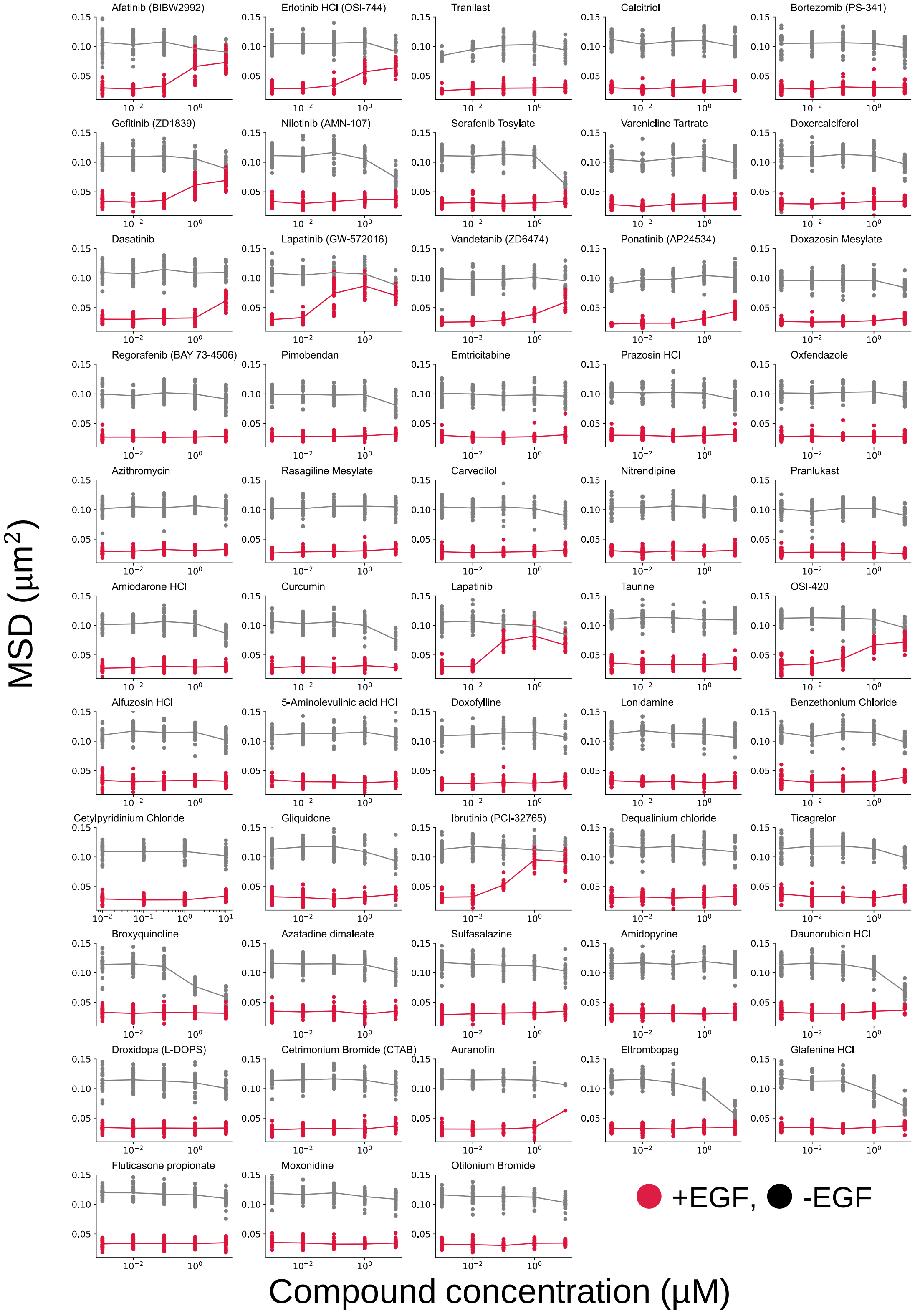


**Supplementary Fig. 4. Compound dose-dependent mobility of EGFR.**

MSDs at Δt=167 ms were obtained for cells treated with various concentrations of the 53 hit compounds. Red and black circles denote values with and without EGF, respectively.

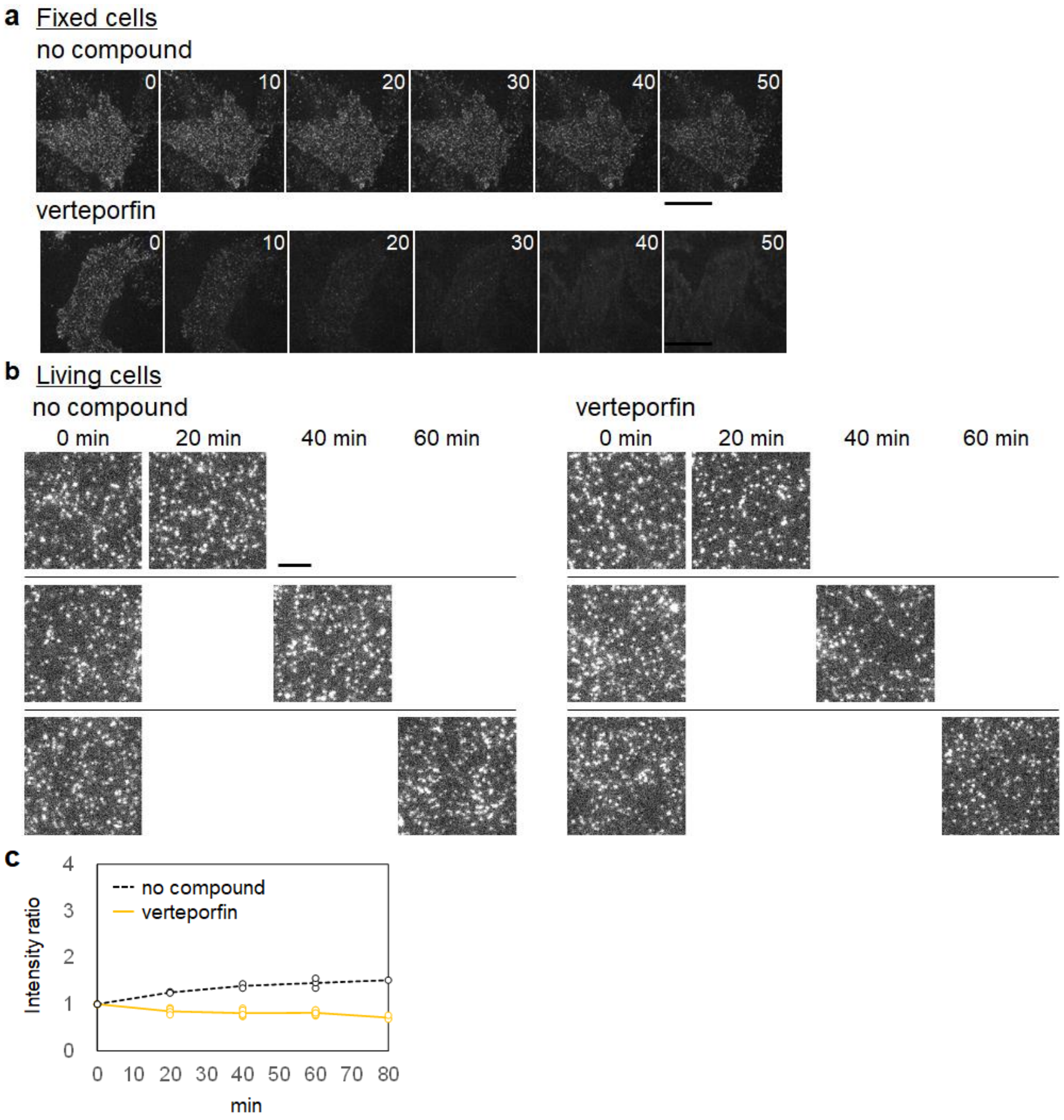


**Supplementary Fig. 5. EGFR internalization and photophysical effects by verteporfin.**

**a,** Timelapse single-molecule images of EGFR-mEGFP in fixed CHO-K1 cells with and without verteporfin treatment. The images were acquired every 10 min after treatment in the same cell. A 488-nm laser repeatedly irradiated the cells, resulting in faster GFP photobleaching due to verteporfin. Scale bar, 5 µm. **b,** Single-molecule images of EGFR-mEGFP in the same living cell with and without verteporfin treatment. The images were acquired at two time points (0 min and 20/40/60 min) after the treatment. Scale bar, 3 µm. **c,** Quantification of the brightness of EGFR-mEGFP on the plasma membrane in the same living cell. The measured intensity at each timepoint was normalized by that at time 0. EGFR-mEGFP on the plasma membrane increased slightly over time under control condition but decreased upon verteporfin treatment, suggesting EGFR internalization by verteporfin.

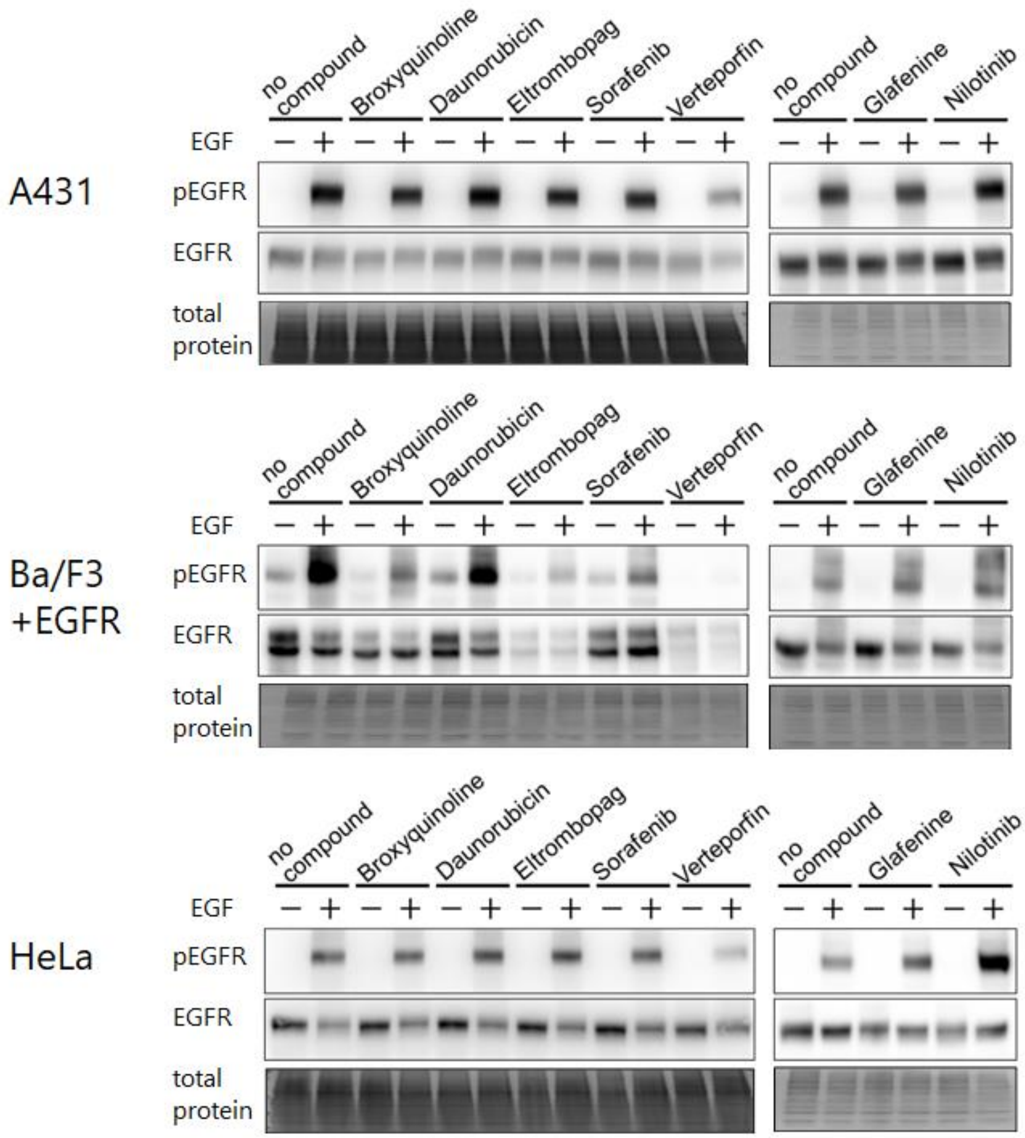


**Supplementary Fig. 6. Effects of non-EGFR-TKI compounds on EGF-induced phosphorylation of EGFR in various cell lines.**

The phosphorylation of EGFR was analyzed in A431, EGFR-expressing Ba/F3, and HeLa cells. After treatment with the indicated compounds for one hour, the cells were stimulated with 60 nM EGF for 2 minutes. EGF stimulation induced the phosphorylation of EGFR in these cell lines even with the application of non-EGFR-TKI compounds, although the extent of the phosphorylation varied with the cell type and compound. In particular, verteporfin-treated cells exhibited defective EGF-induced EGFR phosphorylation, consistent with the inhibition of cluster formation because the EGFR dimer is important for autophosphorylation.

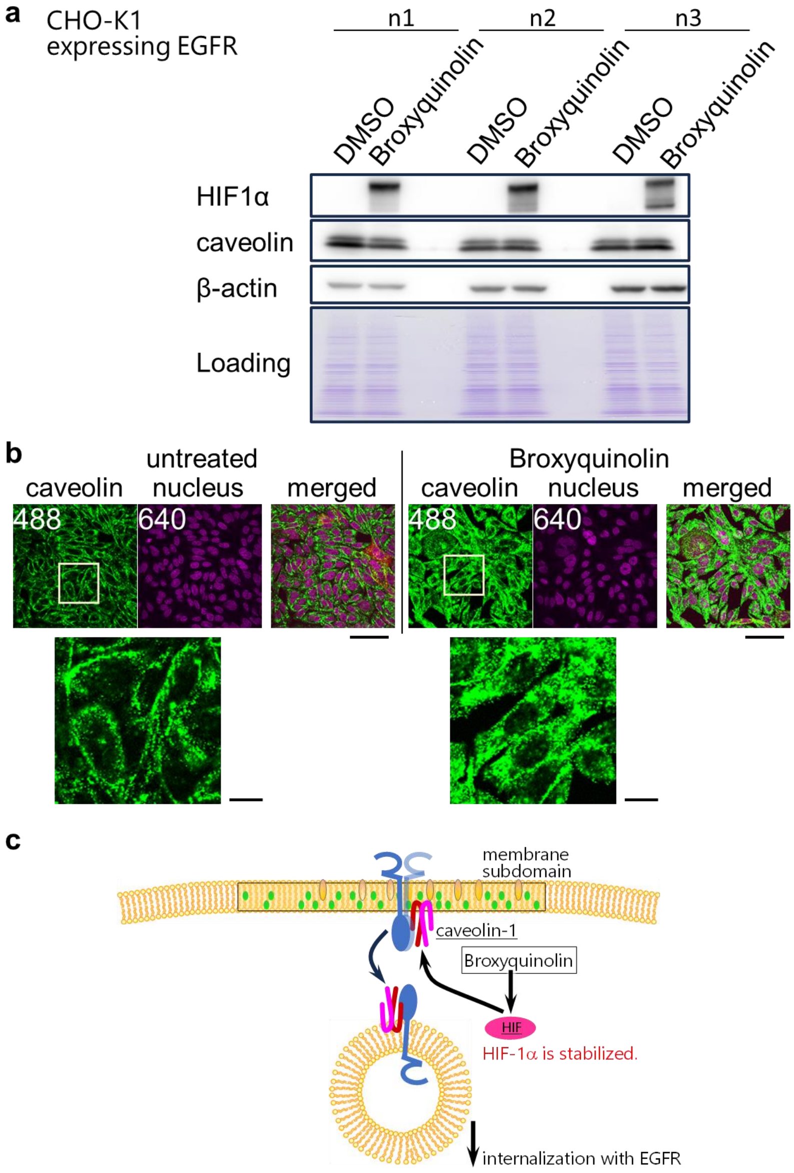


**Supplementary Fig. 7. HIF-1α increase by the broxyquinoline treatment.**

**a,** Western blotting of HIF-1α and caveolin in broxyquinoline-treated and -untreated cells. Results obtained from three replicated experiments are shown. **b,** Immuno-fluorescent images of caveolin in cells with (right panels) and without (left panels) broxyquinoline treatment. Caveolin was stained with Alexa 488-labeled antibody. Fluorescently labeled nuclei were observed with a 640-nm laser. Scale bar, 100 μm. Lower panels represent magnified images of the boxed regions in the upper panels. Scale bars, 20 μm. **c**, A model of the broxyquinoline effect, including HIF-1α production, which induces caveolin internalization with EGFR, based on the current and previous studies[42, 43].

**Supplementary Table 1. Suitability of our method for drug screening.**

All values obtained with our system satisfied the allowance.

| Evaluation | Formula | Our system | Allowance |
|---|---|---|---|
| CV value | $SD/Avg$ | 4% | ≤ 10% |
| S/B ratio | $Avg_{100\%}/Avg_{0\%}$ | 2.7% | ≥ 2% |
| Z'-factor | $Z' = 1 - \frac{(3 \times SD_{positive}) + (3 \times SD_{negative})}{Avg_{positive} - Avg_{negative}}$ | 0.64 | ≥ 0.50 |

### Supplementary Table 2. Number of cells or experiments included in the analysis.

| | | measurement | condition | | | | number | |
|---|---|---|---|---|---|---|---|---|
| **Fig. 1** | **c** | mobility | — | — | EGF | 0 and 60 nM | 900 cells | for each concentration |
| | | clustering | — | — | EGF | 0 and 60 nM | 900 cells | for each concentration |
| | **d** | mobiity | — | — | EGF | 0 and 300 nM | 60 cells | for each concentration |
| | | | gifitinib | 10 µM | EGF | 0 and 300 nM | 60 cells | for each concentration |
| | **e** | mobility | — | — | EGF | 0 - 300 nM | 140 cells | for each concentration |
| | | phosphorylation | — | — | EGF | 0 - 300 nM | 13 experiments | |
| | **f** | mobility | gifitinib | 0.001 – 10 µM | EGF | 0 - 300 nM | 33 cells | for each concentration |
| **Fig. 2** | **a** | mobility | gefitinib | 10 µM | EGF | 60 nM | 10 cells | |
| | **c** | mobility | each compound | 0.001 – 10 µM | — | — | 20 cells | for each concentration |
| | | | | 0.001 – 10 µM | EGF | 60 nM | 20 cells | for each concentration |
| | **d** | mobility | afatinib | 0.001 µM | EGF | 0 and 60 nM | 30 cells (-EGF), 30 cells (+EGF) | |
| | | | | 0.01 µM | | 0 and 60 nM | 30 cells (-EGF), 30 cells (+EGF) | |
| | | | | 0.1 µM | | 0 and 60 nM | 30 cells (-EGF), 30 cells (+EGF) | |
| | | | | 1 µM | | 0 and 60 nM | 30 cells (-EGF), 29 cells (+EGF) | |
| | | | | 10 µM | | 0 and 60 nM | 30 cells (-EGF), 29 cells (+EGF) | |
| | | | elrotinib | 0.001 µM | EGF | 0 and 60 nM | 30 cells (-EGF), 29 cells (+EGF) | |
| | | | | 0.01 µM | | 0 and 60 nM | 30 cells (-EGF), 30 cells (+EGF) | |
| | | | | 0.1 µM | | 0 and 60 nM | 30 cells (-EGF), 29 cells (+EGF) | |
| | | | | 1 µM | | 0 and 60 nM | 30 cells (-EGF), 30 cells (+EGF) | |
| | | | | 10 µM | | 0 and 60 nM | 30 cells (-EGF), 29 cells (+EGF) | |
| | | | OSI-420 | 0.001 µM | EGF | 0 and 60 nM | 28 cells (-EGF), 30 cells (+EGF) | |
| | | | | 0.01 µM | | 0 and 60 nM | 27 cells (-EGF), 28 cells (+EGF) | |
| | | | | 0.1 µM | | 0 and 60 nM | 30 cells (-EGF), 30 cells (+EGF) | |
| | | | | 1 µM | | 0 and 60 nM | 29 cells (-EGF), 29 cells (+EGF) | |
| | | | | 10 µM | | 0 and 60 nM | 23 cells (-EGF), 23 cells (+EGF) | |
| | | | lapatinib | 0.001 µM | EGF | 0 and 60 nM | 30 cells (-EGF), 30 cells (+EGF) | |
| | | | | 0.01 µM | | 0 and 60 nM | 24 cells (-EGF), 25 cells (+EGF) | |
| | | | | 0.1 µM | | 0 and 60 nM | 30 cells (-EGF), 30 cells (+EGF) | |
| | | | | 1 µM | | 0 and 60 nM | 30 cells (-EGF), 30 cells (+EGF) | |
| | | | | 10 µM | | 0 and 60 nM | 30 cells (-EGF), 30 cells (+EGF) | |
| | | | lapatinib ditosylate | 0.001 µM | EGF | 0 and 60 nM | 30 cells (-EGF), 28 cells (+EGF) | |
| | | | | 0.01 µM | | 0 and 60 nM | 30 cells (-EGF), 30 cells (+EGF) | |
| | | | | 0.1 µM | | 0 and 60 nM | 30 cells (-EGF), 28 cells (+EGF) | |
| | | | | 1 µM | | 0 and 60 nM | 29 cells (-EGF), 30 cells (+EGF) | |
| | | | | 10 µM | | 0 and 60 nM | 30 cells (-EGF), 30 cells (+EGF) | |
| | | | ponatinib | 0.001 µM | EGF | 0 and 60 nM | 10 cells (-EGF), 10 cells (+EGF) | |
| | | | | 0.01 µM | | 0 and 60 nM | 30 cells (-EGF), 28 cells (+EGF) | |
| | | | | 0.1 µM | | 0 and 60 nM | 30 cells (-EGF), 29 cells (+EGF) | |
| | | | | 1 µM | | 0 and 60 nM | 30 cells (-EGF), 30 cells (+EGF) | |
| | | | | 10 µM | | 0 and 60 nM | 30 cells (-EGF), 30 cells (+EGF) | |
| | | | vandetanib | 0.001 µM | EGF | 0 and 60 nM | 30 cells (-EGF), 30 cells (+EGF) | |
| | | | | 0.01 µM | | 0 and 60 nM | 30 cells (-EGF), 30 cells (+EGF) | |
| | | | | 0.1 µM | | 0 and 60 nM | 30 cells (-EGF), 30 cells (+EGF) | |
| | | | | 1 µM | | 0 and 60 nM | 30 cells (-EGF), 30 cells (+EGF) | |
| | | | | 10 µM | | 0 and 60 nM | 30 cells (-EGF), 30 cells (+EGF) | |
| | | | dasatinib | 0.001 µM | EGF | 0 and 60 nM | 30 cells (-EGF), 30 cells (+EGF) | |
| | | | | 0.01 µM | | 0 and 60 nM | 30 cells (-EGF), 30 cells (+EGF) | |
| | | | | 0.1 µM | | 0 and 60 nM | 30 cells (-EGF), 30 cells (+EGF) | |
| | | | | 1 µM | | 0 and 60 nM | 30 cells (-EGF), 29 cells (+EGF) | |
| | | | | 10 µM | | 0 and 60 nM | 30 cells (-EGF), 30 cells (+EGF) | |
| | | | iburtinib | 0.001 µM | EGF | 0 and 60 nM | 30 cells (-EGF), 30 cells (+EGF) | |
| | | | | 0.01 µM | | 0 and 60 nM | 30 cells (-EGF), 30 cells (+EGF) | |
| | | | | 0.1 µM | | 0 and 60 nM | 30 cells (-EGF), 29 cells (+EGF) | |
| | | | | 1 µM | | 0 and 60 nM | 30 cells (-EGF), 29 cells (+EGF) | |
| | | | | 10 µM | | 0 and 60 nM | 30 cells (-EGF), 30 cells (+EGF) | |

| | | | | | | | | |
|---|---|---|---|---|---|---|---|---|
| | **e** | mobility | broxyquinoline | 0.001 μM | EGF | 0 and 60 nM | 29 cells (-EGF), 30 cells (+EGF) | |
| | | | | 0.01 μM | | 0 and 60 nM | 28 cells (-EGF), 29 cells (+EGF) | |
| | | | | 0.1 μM | | 0 and 60 nM | 30 cells (-EGF), 30 cells (+EGF) | |
| | | | | 1 μM | | 0 and 60 nM | 24 cells (-EGF), 24 cells (+EGF) | |
| | | | | 10 μM | | 0 and 60 nM | 29 cells (-EGF), 30 cells (+EGF) | |
| | | | daunorubicin | 0.001 μM | EGF | 0 and 60 nM | 30 cells (-EGF), 30 cells (+EGF) | |
| | | | | 0.01 μM | | 0 and 60 nM | 29 cells (-EGF), 29 cells (+EGF) | |
| | | | | 0.1 μM | | 0 and 60 nM | 29 cells (-EGF), 29 cells (+EGF) | |
| | | | | 1 μM | | 0 and 60 nM | 27 cells (-EGF), 20 cells (+EGF) | |
| | | | | 10 μM | | 0 and 60 nM | 12 cells (-EGF), 12 cells (+EGF) | |
| | | | eltrombopag | 0.001 μM | EGF | 0 and 60 nM | 30 cells (-EGF), 30 cells (+EGF) | |
| | | | | 0.01 μM | | 0 and 60 nM | 30 cells (-EGF), 30 cells (+EGF) | |
| | | | | 0.1 μM | | 0 and 60 nM | 30 cells (-EGF), 30 cells (+EGF) | |
| | | | | 1 μM | | 0 and 60 nM | 30 cells (-EGF), 20 cells (+EGF) | |
| | | | | 10 μM | | 0 and 60 nM | 30 cells (-EGF), 30 cells (+EGF) | |
| | | | sorafenib | 0.001 μM | EGF | 0 and 60 nM | 30 cells (-EGF), 30 cells (+EGF) | |
| | | | | 0.01 μM | | 0 and 60 nM | 30 cells (-EGF), 30 cells (+EGF) | |
| | | | | 0.1 μM | | 0 and 60 nM | 30 cells (-EGF), 30 cells (+EGF) | |
| | | | | 1 μM | | 0 and 60 nM | 30 cells (-EGF), 30 cells (+EGF) | |
| | | | | 10 μM | | 0 and 60 nM | 29 cells (-EGF), 29 cells (+EGF) | |
| | | | glafenine | 0.001 μM | EGF | 0 and 60 nM | 30 cells (-EGF), 30 cells (+EGF) | |
| | | | | 0.01 μM | | 0 and 60 nM | 30 cells (-EGF), 29 cells (+EGF) | |
| | | | | 0.1 μM | | 0 and 60 nM | 30 cells (-EGF), 30 cells (+EGF) | |
| | | | | 1 μM | | 0 and 60 nM | 30 cells (-EGF), 30 cells (+EGF) | |
| | | | | 10 μM | | 0 and 60 nM | 30 cells (-EGF), 30 cells (+EGF) | |
| | | | nilotinib | 0.001 μM | EGF | 0 and 60 nM | 30 cells (-EGF), 30 cells (+EGF) | |
| | | | | 0.01 μM | | 0 and 60 nM | 30 cells (-EGF), 30 cells (+EGF) | |
| | | | | 0.1 μM | | 0 and 60 nM | 30 cells (-EGF), 30 cells (+EGF) | |
| | | | | 1 μM | | 0 and 60 nM | 30 cells (-EGF), 29 cells (+EGF) | |
| | | | | 10 μM | | 0 and 60 nM | 30 cells (-EGF), 30 cells (+EGF) | |
| | | | curcumin | 0.001 μM | EGF | 0 and 60 nM | 30 cells (-EGF), 30 cells (+EGF) | |
| | | | | 0.01 μM | | 0 and 60 nM | 30 cells (-EGF), 30 cells (+EGF) | |
| | | | | 0.1 μM | | 0 and 60 nM | 30 cells (-EGF), 30 cells (+EGF) | |
| | | | | 1 μM | | 0 and 60 nM | 30 cells (-EGF), 30 cells (+EGF) | |
| | | | | 10 μM | | 0 and 60 nM | 10 cells (-EGF), 10 cells (+EGF) | |
| **Fig. 3** | **b** | clustering | — | — | EGF | 0.03 nM | 116 cells | |
| | | | | — | | 0.3 nM | 116 cells | |
| | | | | — | | 3 nM | 118 cells | |
| | | | | — | | 30 nM | 116 cells | |
| | | | | — | | 300 nM | 116 cells | |
| | **c** | clustering | each compund | 0 μM, 10 μM | — | — | 20 cells | for each concentration |
| | **e** | clustering | verteporfin | 0.001 μM | — | — | 195 cells | |
| | | | | 0.01 μM | | — | 191 cells | |
| | | | | 0.1 μM | | — | 169 cells | |
| | | | | 1 μM | | — | 115 cells | |
| | | | | 10 μM | | — | 102 cells | |
| | **f** | clustering | — | — | EGF | 0 - 300 nM | 115 cells | for each concentration |
| | | | verteporfin | 0 – 10 μM | | — | 115 cells | for each concentration |
| | **g** | mobility | verteporfin | 0.001 μM | | — | 132 cells | |
| | | | | 0.01 μM | | — | 147 cells | |
| | | | | 0.1 μM | | — | 189 cells | |
| | | | | 1 μM | | — | 192 cells | |
| | | | | 10 μM | | — | 195 cells | |
| **Fig. 4** | **a** | mobility and | — | — | EGF | 0 and 60 nM | 24 cells | for each concentration |
| | | clustering | eltrombopag | 10 μM | EGF | 0 and 60 nM | 30 cells | for each concentration |
| | | | verteporfin | 1 μM | — | — | 39 cells | |
| | | | | 1 μM | EGF | 60 nM | 32 cells | |
| | **g** | brightness | — | — | — | — | 46 cells | |
| | | | broxyquinoline | 10 μM | — | — | 56 cells | |
| | | | daunorubicin | 10 μM | — | — | 61 cells | |
| | | | eltrombopag | 5 μM | — | — | 72 cells | |
| | | | sorafenib | 10 μM | — | — | 47 cells | |
| | | | verteporfin | 1 μM | — | — | 63 cells | |
| | | | glafenine | 10 μM | — | — | 42 cells | |
| | | | nilotinib | 10 μM | — | — | 46 cells | |

**Supplementary Table 3. Compounds in the FDA-approved library used in our screening.**

Compound numbers used in Fig. 2c, name, category of target molecule, and obtained MSD ratios are indicated for every compound.

| # | Name | Target | Relative MSD ratio (fold) |
|---|---|---|---|
| 1 | Phenoxybenzamine | membrane receptor | 0.75 |
| 2 | Pramipexole | membrane receptor | 0.6 |
| 3 | Formoterol | membrane receptor | 0.63 |
| 4 | Mirtazapine | membrane receptor | 0.83 |
| 5 | Fesoterodine | membrane receptor | 0.81 |
| 6 | Ritodrine | membrane receptor | 0.72 |
| 7 | Conivaptan | membrane receptor | 0.71 |
| 8 | Dronedarone | membrane receptor | 0.86 |
| 9 | Dopamine | membrane receptor | 0.87 |
| 10 | Clemastine | membrane receptor | 0.86 |
| 11 | Xylazine | membrane receptor | 0.88 |
| 12 | Valsartan | membrane receptor | 0.7 |
| 13 | Erlotinib | membrane receptor | 3.86 |
| 14 | Epinephrine Bitartrate | membrane receptor | 0.91 |
| 15 | Atropine | membrane receptor | 0.79 |
| 16 | Ketotifen Fumarate | membrane receptor | 0.82 |
| 17 | Diphenhydramine | membrane receptor | 0.41 |
| 18 | Adrenaline | membrane receptor | 0.91 |
| 19 | Naftopidil | membrane receptor | 0.73 |
| 20 | Metoprolol | membrane receptor | 0.55 |
| 21 | Sotalol | membrane receptor | 0.79 |
| 22 | Nizatidine | membrane receptor | 0.82 |
| 23 | Aspartame | membrane receptor | 0.87 |
| 24 | Maraviroc | membrane receptor | 0.79 |
| 25 | Salbutamol | membrane receptor | 0.75 |
| 26 | Candesartan | membrane receptor | ND |
| 27 | Adrenaline | membrane receptor | 0.74 |
| 28 | Phentolamine | membrane receptor | 0.82 |
| 29 | Naphazoline | membrane receptor | 0.83 |
| 30 | Urapidil | membrane receptor | 0.91 |
| 31 | Scopolamine | membrane receptor | 0.89 |
| 32 | Levobetaxolol | membrane receptor | 1.01 |
| 33 | Tiotropium | membrane receptor | 1.08 |
| 34 | Misoprostol | membrane receptor | 1.12 |
| 35 | Metaproterenol Sulfate | membrane receptor | 0.92 |
| 36 | Mesoridazine | membrane receptor | 0.66 |
| 37 | Isoetharine | membrane receptor | 0.45 |
| 38 | Prochlorperazine | membrane receptor | 0.84 |
| 39 | Pindolol | membrane receptor | 0.74 |
| 40 | Diphenylpyraline | membrane receptor | 0.85 |

| | | | |
|---|---|---|---|
| 41 | Lofexidine | membrane receptor | 0.9 |
| 42 | Eltrombopag | membrane receptor | 1.8 |
| 43 | Citrate | membrane receptor | 0.7 |
| 44 | Anisotropine | membrane receptor | 0.8 |
| 45 | Pimozide | membrane receptor | 0.88 |
| 46 | Azelastine | membrane receptor | 0.87 |
| 47 | Pyrilamine | membrane receptor | 0.85 |
| 48 | Serotonin | membrane receptor | 0.8 |
| 49 | Doxylamine | membrane receptor | 0.83 |
| 50 | Pipenzolate | membrane receptor | 0.85 |
| 51 | Pilocarpine | membrane receptor | 0.87 |
| 52 | Bromocriptine | membrane receptor | 0.91 |
| 53 | Plerixafor | membrane receptor | 0.9 |
| 54 | Methoxamine | membrane receptor | 0.98 |
| 55 | Nalmefene | membrane receptor | 0.9 |
| 56 | Otilonium | membrane receptor | 1.27 |
| 57 | Aceclidine | membrane receptor | 1.04 |
| 58 | Mepenzolate | membrane receptor | 0.97 |
| 59 | Thioridazine | membrane receptor | 0.87 |
| 60 | Dicyclomine | membrane receptor | 1.01 |
| 61 | Triflupromazine | membrane receptor | 0.92 |
| 62 | Metaraminol | membrane receptor | 0.86 |
| 63 | Pheniramine | membrane receptor | 1.03 |
| 64 | Tolvaptan | membrane receptor | 1.06 |
| 65 | Moxonidine | membrane receptor | 1.32 |
| 66 | Carbachol | membrane receptor | 0.99 |
| 67 | Bismuth | membrane receptor | 0.93 |
| 68 | Benztropine | membrane receptor | 0.92 |
| 69 | Terfenadine | membrane receptor | 1.08 |
| 70 | Ractopamine | membrane receptor | 1.06 |
| 71 | Apomorphine | membrane receptor | 1.21 |
| 72 | Procyclidine | membrane receptor | 1.07 |
| 73 | Almotriptan | membrane receptor | 0.85 |
| 74 | Oxprenolol | membrane receptor | 0.95 |
| 75 | Hyoscyamine | membrane receptor | 0.98 |
| 76 | Desloratadine | membrane receptor | 0.65 |
| 77 | Acebutolol | membrane receptor | 0.91 |
| 78 | Mirabegron | membrane receptor | 0.79 |
| 79 | Rimonabant | membrane receptor | 1.06 |
| 80 | Cimetidine | membrane receptor | 0.67 |
| 81 | Betahistine | membrane receptor | 0.88 |
| 82 | Pazopanib | membrane receptor | 1.06 |
| 83 | Ticagrelor | membrane receptor | 1.71 |
| 84 | Propranolol | membrane receptor | 0.81 |
| 85 | Ipratropium | membrane receptor | 0.94 |
| 86 | Guanabenz Acetate | membrane receptor | 0.9 |
| 87 | Noradrenaline | membrane receptor | 0.6 |

| | | | |
|---|---|---|---|
| 88 | Tripelennamine | membrane receptor | 1.1 |
| 89 | Choline Chloride | membrane receptor | 1.0 |
| 90 | Darifenacin | membrane receptor | 1.0 |
| 91 | Doxofylline | membrane receptor | 1.25 |
| 92 | Orphenadrine | membrane receptor | 1.0 |
| 93 | Haloperidol | membrane receptor | 0.94 |
| 94 | Tropicamide | membrane receptor | 1.11 |
| 95 | Tolterodine | membrane receptor | 0.95 |
| 96 | Trospium | membrane receptor | 0.96 |
| 97 | Alfuzosin | membrane receptor | 1.31 |
| 98 | Loxapine Succinate | membrane receptor | 1.03 |
| 99 | Cyproheptadine | membrane receptor | 0.9 |
| 100 | Bismuth | membrane receptor | 1.05 |
| 101 | Azatadine | membrane receptor | 1.25 |
| 102 | Medetomidine | membrane receptor | 1.12 |
| 103 | Histamine | membrane receptor | 0.94 |
| 104 | Azilsartan | membrane receptor | 0.91 |
| 105 | Solifenacin | membrane receptor | 1.05 |
| 106 | Pergolide | membrane receptor | 1.15 |
| 107 | Montelukast | membrane receptor | 1.23 |
| 108 | Fexofenadine | membrane receptor | 1.03 |
| 109 | Meptazinol | membrane receptor | 0.97 |
| 110 | Azilsartan | membrane receptor | 1.07 |
| 111 | Lurasidone | membrane receptor | 0.86 |
| 112 | Eprosartan | membrane receptor | 1.14 |
| 113 | Droxidopa | membrane receptor | 1.29 |
| 114 | Esmolol | membrane receptor | 1.01 |
| 115 | Fosaprepitant | membrane receptor | 0.86 |
| 116 | Bepotastine | membrane receptor | 1.12 |
| 117 | Droperidol | membrane receptor | 1.19 |
| 118 | Trifluoperazine | membrane receptor | 0.86 |
| 119 | Ropinirole | membrane receptor | 0.93 |
| 120 | Antazoline | membrane receptor | 0.9 |
| 121 | Adrenalone | membrane receptor | 0.91 |
| 122 | Blonanserin | membrane receptor | 0.6 |
| 123 | Afatinib | membrane receptor | 2.63 |
| 124 | Lapatinib | membrane receptor | 3.8 |
| 125 | Oxybutynin | membrane receptor | 0.87 |
| 126 | Imatinib | membrane receptor | 1.21 |
| 127 | Aminophylline | membrane receptor | 1.01 |
| 128 | Famotidine | membrane receptor | 0.91 |
| 129 | Losartan | membrane receptor | 0.98 |
| 130 | Ramelteon | membrane receptor | 0.86 |
| 131 | Biperiden | membrane receptor | 0.93 |
| 132 | Loperamide | membrane receptor | 1.03 |
| 133 | Methyldopa | membrane receptor | 0.67 |
| 134 | Gallamine | membrane receptor | 0.93 |

| | | | |
|---|---|---|---|
| 135 | Cinacalcet | membrane receptor | 0.54 |
| 136 | Asenapine | membrane receptor | 0.91 |
| 137 | Aripiprazole | membrane receptor | 1.06 |
| 138 | Ambrisentan | membrane receptor | 1.03 |
| 139 | Naratriptan | membrane receptor | 1.09 |
| 140 | Cetirizine | membrane receptor | 1.12 |
| 141 | Zolmitriptan | membrane receptor | 0.79 |
| 142 | Pranlukast | membrane receptor | 1.37 |
| 143 | Nebivolol | membrane receptor | 1.02 |
| 144 | Carvedilol | membrane receptor | 1.25 |
| 145 | Meglumine | membrane receptor | 0.95 |
| 146 | Bethanechol | membrane receptor | 0.76 |
| 147 | Prasugrel | membrane receptor | 1.12 |
| 148 | Chlorpromazine | membrane receptor | 0.69 |
| 149 | Pramipexole | membrane receptor | 0.88 |
| 150 | Masitinib | membrane receptor | 1.07 |
| 151 | Iloperidone | membrane receptor | 0.85 |
| 152 | Vandetanib | membrane receptor | 1.67 |
| 153 | Ranitidine | membrane receptor | 0.63 |
| 154 | Crizotinib | membrane receptor | 1.08 |
| 155 | Clozapine | membrane receptor | 0.71 |
| 156 | Vismodegib | membrane receptor | 0.55 |
| 157 | Sunitinib | membrane receptor | ND |
| 158 | Cabozantinib | membrane receptor | 1.22 |
| 159 | Bimatoprost | membrane receptor | 1.0 |
| 160 | Acetylcholine | membrane receptor | 0.76 |
| 161 | Clonidine | membrane receptor | 0.65 |
| 162 | Trimebutine | membrane receptor | 0.89 |
| 163 | Axitinib | membrane receptor | 1.16 |
| 164 | Cilostazol | membrane receptor | 0.96 |
| 165 | Dexmedetomidine | membrane receptor | 1.03 |
| 166 | Betaxolol | membrane receptor | 0.85 |
| 167 | Chlorpheniramine | membrane receptor | 0.79 |
| 168 | Detomidine | membrane receptor | 0.97 |
| 169 | Aprepitant | membrane receptor | 1.03 |
| 170 | Naltrexone | membrane receptor | 1.03 |
| 171 | Levosulpiride | membrane receptor | 0.97 |
| 172 | Betaxolol hydrochloride | membrane receptor | 1.1 |
| 173 | Prazosin | membrane receptor | 1.38 |
| 174 | Lapatinib | membrane receptor | 2.67 |
| 175 | Imatinib Mesylate | membrane receptor | 0.99 |
| 176 | Adenine | membrane receptor | 1.06 |
| 177 | Quetiapine | membrane receptor | 0.91 |
| 178 | Ziprasidone | membrane receptor | 1.03 |
| 179 | Gefitinib | membrane receptor | 2.82 |
| 180 | Ticlopidine | membrane receptor | 1.01 |
| 181 | Erlotinib | membrane receptor | 2.44 |

| | | | |
|---|---|---|---|
| 182 | Olanzapine | membrane receptor | 1.07 |
| 183 | Adenine | membrane receptor | 1.14 |
| 184 | Olopatadine | membrane receptor | 0.65 |
| 185 | Dasatinib | membrane receptor | 2.4 |
| 186 | Zafirlukast | membrane receptor | 0.85 |
| 187 | Doxazosin | membrane receptor | 1.29 |
| 188 | Sumatriptan | membrane receptor | 0.96 |
| 189 | Meclizine | membrane receptor | 0.67 |
| 190 | Agomelatine | membrane receptor | 0.96 |
| 191 | Oxymetazoline | membrane receptor | 0.8 |
| 192 | Chlorprothixene | membrane receptor | 0.96 |
| 193 | Alprostadil | membrane receptor | 0.84 |
| 194 | Irbesartan | membrane receptor | 0.82 |
| 195 | Adenine | membrane receptor | 1.08 |
| 196 | Domperidone | membrane receptor | 0.97 |
| 197 | Mizolastine | membrane receptor | 1.05 |
| 198 | Dyphylline | membrane receptor | 0.95 |
| 199 | Pazopanib | membrane receptor | 1.02 |
| 200 | Amisulpride | membrane receptor | 0.98 |
| 201 | Amiodarone | membrane receptor | 1.31 |
| 202 | Tizanidine | membrane receptor | 0.98 |
| 203 | Clopidogrel | membrane receptor | 0.81 |
| 204 | Rocuronium | membrane receptor | 0.87 |
| 205 | Methscopolamine | membrane receptor | 0.86 |
| 206 | Nilotinib | membrane receptor | 1.86 |
| 207 | Adenosine | membrane receptor | 0.89 |
| 208 | Lonidamine | enzyme | 1.42 |
| 209 | Uridine | enzyme | 0.59 |
| 210 | Methimazole | enzyme | 0.81 |
| 211 | Diclofenac | enzyme | 0.97 |
| 212 | Ibandronate | enzyme | 0.93 |
| 213 | Ketorolac | enzyme | 0.7 |
| 214 | Acarbose | enzyme | 0.7 |
| 215 | Ketoprofen | enzyme | 0.64 |
| 216 | Uracil | enzyme | 0.84 |
| 217 | Celecoxib | enzyme | 0.78 |
| 218 | Moclobemide | enzyme | 1.03 |
| 219 | Ibuprofen | enzyme | 1.02 |
| 220 | Carfilzomib | enzyme | 0.94 |
| 221 | Cobicistat | enzyme | 0.76 |
| 222 | Diclofenac | enzyme | 0.8 |
| 223 | Etodolac | enzyme | 0.88 |
| 224 | Ampiroxicam | enzyme | 0.94 |
| 225 | Rosuvastatin | enzyme | 1.03 |
| 226 | Dichlorphenamide | enzyme | 1.14 |
| 227 | Rasagiline | enzyme | 1.43 |
| 228 | Benzydamine | enzyme | 1.01 |

| | | | |
|---|---|---|---|
| 229 | Anisindione | enzyme | 0.85 |
| 230 | Perindopril | enzyme | 0.85 |
| 231 | Gemcitabine | enzyme | 0.93 |
| 232 | Sulindac | enzyme | 0.9 |
| 233 | Temocapril | enzyme | 0.99 |
| 234 | Sildenafil | enzyme | 0.76 |
| 235 | Capecitabine | enzyme | 1.0 |
| 236 | Tenoxicam | enzyme | 0.77 |
| 237 | Sodium salicylate | enzyme | 1.01 |
| 238 | Tadalafil | enzyme | 0.92 |
| 239 | Rolipram | enzyme | 0.83 |
| 240 | Cyclosporine | enzyme | 0.86 |
| 241 | Vardenafil | enzyme | 0.92 |
| 242 | Oxaprozin | enzyme | 1.0 |
| 243 | Methylthiouracil | enzyme | 1.03 |
| 244 | Mitoxantrone | enzyme | 0.9 |
| 245 | Risedronate | enzyme | 0.84 |
| 246 | Rofecoxib | enzyme | 1.14 |
| 247 | Roflumilast | enzyme | 0.45 |
| 248 | Leflunomide | enzyme | 0.92 |
| 249 | Allopurinol | enzyme | 0.92 |
| 250 | Zaltoprofen | enzyme | 0.82 |
| 251 | Irinotecan | enzyme | 0.87 |
| 252 | Dipyridamole | enzyme | ND |
| 253 | Linagliptin | enzyme | 0.89 |
| 254 | Topotecan | enzyme | 0.97 |
| 255 | Orlistat | enzyme | 0.84 |
| 256 | Fenoprofen | enzyme | 0.97 |
| 257 | Ibuprofen | enzyme | 1.0 |
| 258 | Nepafenac | enzyme | 1.0 |
| 259 | Bufexamac | enzyme | 0.86 |
| 260 | Rivastigmine | enzyme | 0.85 |
| 261 | Pitavastatin | enzyme | 1.19 |
| 262 | Hydroxyurea | enzyme | 0.88 |
| 263 | Anagrelide | enzyme | 1.1 |
| 264 | Esomeprazole | enzyme | 1.02 |
| 265 | Carbidopa | enzyme | 0.84 |
| 266 | Fosinopril | enzyme | 0.81 |
| 267 | Triflusal | enzyme | 0.88 |
| 268 | Finasteride | enzyme | 1.22 |
| 269 | Pimobendan | enzyme | 1.55 |
| 270 | Irinotecan | enzyme | 0.91 |
| 271 | Cladribine | enzyme | 0.99 |
| 272 | Voglibose | enzyme | 0.94 |
| 273 | Dabigatran | enzyme | 0.82 |
| 274 | Rivaroxaban | enzyme | 0.89 |
| 275 | Aliskiren | enzyme | 0.73 |

| | | | |
|---|---|---|---|
| 276 | TAME | enzyme | 0.92 |
| 277 | Dexlansoprazole | enzyme | 1.09 |
| 278 | Alendronate | enzyme | 0.93 |
| 279 | Pranoprofen | enzyme | 0.8 |
| 280 | Pravastatin | enzyme | 0.95 |
| 281 | Mefenamic | enzyme | 0.65 |
| 282 | Naproxen | enzyme | 0.95 |
| 283 | Daunorubicin | enzyme | 1.99 |
| 284 | Tolfenamic | enzyme | 0.77 |
| 285 | Exemestane | enzyme | 1.0 |
| 286 | Tranexamic | enzyme | 0.81 |
| 287 | Gabexate | enzyme | 0.9 |
| 288 | Mofetil | enzyme | 0.84 |
| 289 | Aspirin | enzyme | 1.01 |
| 290 | Benazepril | enzyme | 0.86 |
| 291 | Phenylbutazone | enzyme | 0.66 |
| 292 | Imidapril | enzyme | 0.93 |
| 293 | Cytidine | enzyme | 1.05 |
| 294 | Enalaprilat | enzyme | 0.91 |
| 295 | Tacrine | enzyme | 1.0 |
| 296 | Simvastatin | enzyme | 0.98 |
| 297 | Racecadotril | enzyme | 0.7 |
| 298 | Atorvastatin | enzyme | 1.14 |
| 299 | Carmofur | enzyme | 0.99 |
| 300 | Lisinopril | enzyme | 0.82 |
| 301 | Phenindione | enzyme | 1.13 |
| 302 | Neostigmine | enzyme | 0.83 |
| 303 | Avanafil | enzyme | 1.05 |
| 304 | Pemetrexed | enzyme | 1.06 |
| 305 | Nimesulide | enzyme | 0.76 |
| 306 | Ramipril | enzyme | 0.71 |
| 307 | Physostigmine | enzyme | 0.91 |
| 308 | Moexipril | enzyme | 0.74 |
| 309 | Captopril | enzyme | 0.86 |
| 310 | Bortezomib | enzyme | 1.41 |
| 311 | Glycyrrhizinate | enzyme | 0.88 |
| 312 | Dabigatran | enzyme | 1.24 |
| 313 | Cilazapril | enzyme | 0.93 |
| 314 | Enalapril | enzyme | 0.65 |
| 315 | Fluvastatin | enzyme | 1.16 |
| 316 | Propylthiouracil | enzyme | 0.62 |
| 317 | Zileuton | enzyme | 0.97 |
| 318 | Ozagrel | enzyme | 0.64 |
| 319 | Abiraterone | enzyme | 0.89 |
| 320 | Doxifluridine | enzyme | 0.91 |
| 321 | Lornoxicam | enzyme | 0.93 |
| 322 | Donepezil | enzyme | 0.75 |

| | | | |
|---|---|---|---|
| 323 | Esomeprazole | enzyme | 0.83 |
| 324 | Carbenoxolone | enzyme | 0.92 |
| 325 | Tolmetin | enzyme | 1.0 |
| 326 | Physostigmine | enzyme | 0.88 |
| 327 | Amfenac | enzyme | 0.95 |
| 328 | Benserazide | enzyme | 0.91 |
| 329 | Teniposide | enzyme | 0.83 |
| 330 | Floxuridine | enzyme | 0.74 |
| 331 | Valdecoxib | enzyme | 1.01 |
| 332 | Nabumetone | enzyme | 1.04 |
| 333 | Aminocaproic | enzyme | 0.91 |
| 334 | Tegafur | enzyme | 0.77 |
| 335 | Aminoglutethimide | enzyme | 1.06 |
| 336 | Ethoxzolamide | enzyme | 0.8 |
| 337 | Diclofenac | enzyme | 0.75 |
| 338 | Thioguanine | enzyme | 0.82 |
| 339 | Risedronic | enzyme | 0.84 |
| 340 | Mercaptopurine | enzyme | 0.66 |
| 341 | Acemetacin | enzyme | 1.02 |
| 342 | Pamidronate | enzyme | 0.94 |
| 343 | Flurbiprofen | enzyme | 1.02 |
| 344 | Disulfiram | enzyme | 0.61 |
| 345 | Zoledronic | enzyme | 0.87 |
| 346 | Epalrestat | enzyme | 1.12 |
| 347 | Abiraterone | enzyme | 0.67 |
| 348 | Rolipram | enzyme | 0.93 |
| 349 | Carbimazole | enzyme | 1.0 |
| 350 | Brinzolamide | enzyme | 0.92 |
| 351 | Ozagrel | enzyme | 0.83 |
| 352 | Vorinostat | enzyme | 1.01 |
| 353 | Thiouracil | enzyme | 0.85 |
| 354 | Argatroban | enzyme | 0.93 |
| 355 | Pimecrolimus | enzyme | 0.78 |
| 356 | Gimeracil | enzyme | 1.06 |
| 357 | Fludarabine | enzyme | 0.79 |
| 358 | Mycophenolic | enzyme | 0.96 |
| 359 | Tacrolimus | enzyme | 0.88 |
| 360 | Pralatrexate | enzyme | 0.82 |
| 361 | Azacitidine | enzyme | 0.79 |
| 362 | Nialamide | enzyme | 0.98 |
| 363 | Benzoic | ion channel | 1.03 |
| 364 | Amantadine | ion channel | 0.94 |
| 365 | Triamterene | ion channel | 0.8 |
| 366 | Vecuronium | ion channel | 0.98 |
| 367 | Amlodipine | ion channel | 0.92 |
| 368 | Nifedipine | ion channel | 0.74 |
| 369 | Memantine | ion channel | 0.83 |

| | | | |
|---|---|---|---|
| 370 | Rufinamide | ion channel | 0.99 |
| 371 | Articaine | ion channel | 0.81 |
| 372 | Flunarizine | ion channel | 0.91 |
| 373 | Gluconate | ion channel | 0.88 |
| 374 | Clevidipine | ion channel | 0.75 |
| 375 | Hexamethonium | ion channel | 0.82 |
| 376 | Nicardipine | ion channel | 1.08 |
| 377 | Phenytoin | ion channel | 0.9 |
| 378 | Tropisetron | ion channel | 0.77 |
| 379 | Ivabradine | ion channel | 0.85 |
| 380 | Mepivacaine | ion channel | 0.91 |
| 381 | Gabapentin | ion channel | 0.99 |
| 382 | Niflumic | ion channel | 0.72 |
| 383 | Oxybuprocaine | ion channel | 0.94 |
| 384 | Valproic | ion channel | 1.03 |
| 385 | Proparacaine | ion channel | 1.01 |
| 386 | Topiramate | ion channel | 0.81 |
| 387 | Atracurium | ion channel | 0.96 |
| 388 | Disopyramide | ion channel | 0.72 |
| 389 | Quipazine | ion channel | 0.81 |
| 390 | Azasetron | ion channel | 0.84 |
| 391 | Cilnidipine | ion channel | 0.91 |
| 392 | Diltiazem | ion channel | 0.62 |
| 393 | Oxethazaine | ion channel | 0.95 |
| 394 | Nitrendipine | ion channel | 1.27 |
| 395 | Isradipine | ion channel | 0.89 |
| 396 | Dofetilide | ion channel | 0.82 |
| 397 | Bupivacaine | ion channel | 1.01 |
| 398 | Ondansetron | ion channel | 0.75 |
| 399 | Propafenone | ion channel | 0.77 |
| 400 | ATP | ion channel | 0.8 |
| 401 | Phenytoin | ion channel | 1.05 |
| 402 | Lacidipine | ion channel | 1.04 |
| 403 | Ibutilide | ion channel | 0.7 |
| 404 | Zonisamide | ion channel | 0.69 |
| 405 | Pancuronium | ion channel | 0.77 |
| 406 | Varenicline | ion channel | 1.45 |
| 407 | Cisatracurium | ion channel | 0.78 |
| 408 | Etomidate | ion channel | 0.94 |
| 409 | Mexiletine | ion channel | 1.03 |
| 410 | Gabapentin | ion channel | 0.86 |
| 411 | Benidipine | ion channel | 1 |
| 412 | Lomerizine | ion channel | 0.93 |
| 413 | Amiloride | ion channel | 0.68 |
| 414 | Nisoldipine | ion channel | 0.88 |
| 415 | Amlodipine | ion channel | 0.76 |
| 416 | Nimodipine | ion channel | 0.96 |

| | | | |
|---|---|---|---|
| 417 | Nicotinic | ion channel | 0.88 |
| 418 | Lamotrigine | ion channel | 1.04 |
| 419 | Pramoxine | ion channel | 0.92 |
| 420 | Divalproex | ion channel | 1.04 |
| 421 | Felodipine | ion channel | 1.15 |
| 422 | Felbamate | ion channel | 0.96 |
| 423 | Benzocaine | ion channel | 1.14 |
| 424 | Ropivacaine | ion channel | 0.99 |
| 425 | Manidipine | ion channel | 1.07 |
| 426 | Carbamazepine | ion channel | 1.02 |
| 427 | Palonosetron | ion channel | 0.99 |
| 428 | Azelnidipine | ion channel | 0.72 |
| 429 | Flumazenil | ion channel | 1.02 |
| 430 | Vitamin D3 | nuclear receptor | 1.03 |
| 431 | Dichlorisone | nuclear receptor | 0.73 |
| 432 | Toremifene | nuclear receptor | 0.82 |
| 433 | Fluorometholone | nuclear receptor | 0.65 |
| 434 | Ethisterone | nuclear receptor | 1.13 |
| 435 | Fluocinonide | nuclear receptor | 0.98 |
| 436 | Adapalene | nuclear receptor | 0.87 |
| 437 | Flutamide | nuclear receptor | 0.97 |
| 438 | Hydrocortisone | nuclear receptor | 0.91 |
| 439 | Medrysone | nuclear receptor | 0.73 |
| 440 | Betamethasone | nuclear receptor | 1.07 |
| 441 | Ursodiol | nuclear receptor | 0.63 |
| 442 | Desonide | nuclear receptor | 0.97 |
| 443 | Loteprednol | nuclear receptor | 0.91 |
| 444 | Rosiglitazone | nuclear receptor | 0.95 |
| 445 | Ethynodiol | nuclear receptor | 0.95 |
| 446 | Pioglitazone | nuclear receptor | 0.86 |
| 447 | Estradiol valerate | nuclear receptor | 1.09 |
| 448 | Mifepristone | nuclear receptor | 1.1 |
| 449 | Spironolactone | nuclear receptor | 1.03 |
| 450 | Fenofibrate | nuclear receptor | 0.83 |
| 451 | Betamethasone | nuclear receptor | 0.93 |
| 452 | Megestrol | nuclear receptor | 0.88 |
| 453 | Meprednisone | nuclear receptor | 0.93 |
| 454 | Canrenoate | nuclear receptor | 1.08 |
| 455 | Liothyronine | nuclear receptor | 0.96 |
| 456 | Tiratricol | nuclear receptor | 0.8 |
| 457 | Estrone | nuclear receptor | 0.89 |
| 458 | Fluticasone | nuclear receptor | 1.76 |
| 459 | Budesonide | nuclear receptor | 0.92 |
| 460 | Fulvestrant | nuclear receptor | 0.97 |
| 461 | Tamoxifen | nuclear receptor | 0.88 |
| 462 | Bexarotene | nuclear receptor | 1.08 |
| 463 | Dexamethasone | nuclear receptor | 0.86 |

| | | | |
|---|---|---|---|
| 464 | Oxymetholone | nuclear receptor | 0.98 |
| 465 | Tazarotene | nuclear receptor | 1.19 |
| 466 | Flumethasone | nuclear receptor | 1.04 |
| 467 | Bazedoxifene | nuclear receptor | 0.95 |
| 468 | Halobetasol Propionate | nuclear receptor | 1.09 |
| 469 | Diethylstilbestrol | nuclear receptor | 0.82 |
| 470 | Mestranol | nuclear receptor | 0.85 |
| 471 | Dydrogesterone | nuclear receptor | 0.79 |
| 472 | Estradiol | nuclear receptor | 0.85 |
| 473 | Rosiglitazone | nuclear receptor | 0.91 |
| 474 | butyrate | nuclear receptor | 0.71 |
| 475 | Difluprednate | nuclear receptor | 1.02 |
| 476 | Mometasone | nuclear receptor | 0.73 |
| 477 | Triamcinolone | nuclear receptor | 0.89 |
| 478 | Deflazacort | nuclear receptor | 0.7 |
| 479 | Fluocinolone | nuclear receptor | 0.96 |
| 480 | Tretinoin | nuclear receptor | 0.95 |
| 481 | Calcitriol | nuclear receptor | 1.39 |
| 482 | Dexamethasone | nuclear receptor | 1.01 |
| 483 | Doxercalciferol | nuclear receptor | 1.45 |
| 484 | Estriol | nuclear receptor | 0.75 |
| 485 | Altrenogest | nuclear receptor | 0.8 |
| 486 | Betamethasone | nuclear receptor | 0.9 |
| 487 | Calcifediol | nuclear receptor | 1.17 |
| 488 | Alfacalcidol | nuclear receptor | 0.87 |
| 489 | Clofibrate | nuclear receptor | 0.87 |
| 490 | Triamcinolone | nuclear receptor | 0.9 |
| 491 | Nateglinide | transporter | 1.03 |
| 492 | Reboxetine | transporter | 0.81 |
| 493 | Imipramine | transporter | 1.06 |
| 494 | Gliclazide | transporter | 0.91 |
| 495 | Bendroflumethiazide | transporter | 0.86 |
| 496 | Amitriptyline | transporter | 1.04 |
| 497 | Benzthiazide | transporter | 1.06 |
| 498 | Ivacaftor | transporter | ND |
| 499 | Methyclothiazide | transporter | 0.94 |
| 500 | Indapamide | transporter | 0.84 |
| 501 | Venlafaxine | transporter | 1.04 |
| 502 | Sertraline | transporter | 1.06 |
| 503 | Tolbutamide | transporter | 1.01 |
| 504 | Mitiglinide | transporter | 0.91 |
| 505 | Repaglinide | transporter | 1.02 |
| 506 | Dapoxetine | transporter | 0.83 |
| 507 | Gliquidone | transporter | 1.26 |
| 508 | Duloxetine | transporter | 0.74 |
| 509 | Torsemide | transporter | 0.96 |
| 510 | Chlorothiazide | transporter | 0.96 |

| | | | |
|---|---|---|---|
| 511 | Paroxetine | transporter | 0.55 |
| 512 | Trimipramine | transporter | 0.44 |
| 513 | Fluvoxamine | transporter | 0.96 |
| 514 | Trichlormethiazide | transporter | 0.9 |
| 515 | Glipizide | transporter | 0.88 |
| 516 | Benzbromarone | transporter | 1.0 |
| 517 | Clomipramine | transporter | 0.9 |
| 518 | Tolazamide | transporter | 0.96 |
| 519 | Bumetanide | transporter | 0.92 |
| 520 | Amoxapine | transporter | 0.86 |
| 521 | Ezetimibe | transporter | 1.17 |
| 522 | Guanethidine | transporter | 0.84 |
| 523 | Nicorandil | transporter | 1.09 |
| 524 | Nomifensine | transporter | 0.89 |
| 525 | Maprotiline | transporter | 0.78 |
| 526 | Meticrane | transporter | 0.8 |
| 527 | Atomoxetine | transporter | 0.89 |
| 528 | Milnacipran | transporter | 1.01 |
| 529 | Pinacidil | transporter | 0.94 |
| 530 | Chlorpropamide | transporter | 1.07 |
| 531 | Fluoxetine | transporter | 0.93 |
| 532 | Everolimus | non-receptor kinase | 1.07 |
| 533 | Sorafenib | non-receptor kinase | 1.48 |
| 534 | Vemurafenib | non-receptor kinase | 1.0 |
| 535 | Phenformin | non-receptor kinase | 0.94 |
| 536 | Ponatinib | non-receptor kinase | 1.78 |
| 537 | Ibrutinib | non-receptor kinase | 2.17 |
| 538 | Regorafenib | non-receptor kinase | 1.68 |
| 539 | Metformin | non-receptor kinase | 0.75 |
| 540 | Dabrafenib | non-receptor kinase | 0.77 |
| 541 | Temsirolimus | non-receptor kinase | 0.7 |
| 542 | Thalidomide | cytokine | 0.98 |
| 543 | Pomalidomide | cytokine | 1.12 |
| 544 | Lenalidomide | cytokine | 0.76 |
| 545 | Bindarit | cytokine | 1.0 |
| 546 | Sulfanilamide | other | 0.86 |
| 547 | Edaravone | other | 0.88 |
| 548 | Methoxyestradiol | other | 0.94 |
| 549 | Cyclamic | other | 1.07 |
| 550 | Lithocholic | other | 0.87 |
| 551 | Emtricitabine | other | 1.26 |
| 552 | Genistein | other | 0.97 |
| 553 | Valnemulin | other | 1.08 |
| 554 | Monofluorophosphate | other | 0.82 |
| 555 | Ethambutol | other | 1.0 |
| 556 | Leucovorin | other | 0.79 |
| 557 | Gatifloxacin | other | 1.07 |

| | | | |
|---|---|---|---|
| 558 | Temozolomide | other | 0.96 |
| 559 | Sulfasalazine | other | 1.29 |
| 560 | Ouabain | other | 0.64 |
| 561 | Clofibric | other | 0.91 |
| 562 | Nitazoxanide | other | 1 |
| 563 | Azaperone | other | 0.61 |
| 564 | Nithiamide | other | 0.65 |
| 565 | Allylthiourea | other | 0.86 |
| 566 | Cysteamine | other | 0.85 |
| 567 | Zoxazolamine | other | 0.93 |
| 568 | Phenazopyridine | other | 0.72 |
| 569 | Penciclovir | other | 0.95 |
| 570 | Vincristine | other | 0.94 |
| 571 | Clinafloxacin | other | 0.67 |
| 572 | Natamycin | other | 0.93 |
| 573 | Ritonavir | other | 1.04 |
| 574 | Alverine Citrate | other | 0.72 |
| 575 | Didanosine | other | 1.04 |
| 576 | Besifloxacin | other | 0.98 |
| 577 | Amidopyrine | other | 1.28 |
| 578 | Triclabendazole | other | 0.61 |
| 579 | Dicloxacillin | other | 0.98 |
| 580 | Vinorelbine | other | 1.15 |
| 581 | Chlorocresol | other | 1.13 |
| 582 | Telaprevir | other | 0.99 |
| 583 | Isovaleramide | other | 1.04 |
| 584 | Danofloxacin | other | 0.68 |
| 585 | Sulconazole | other | 0.7 |
| 586 | Enrofloxacin | other | 0.68 |
| 587 | Tilmicosin | other | 0.92 |
| 588 | Ethionamide | other | 0.95 |
| 589 | Thiamine | other | 0.91 |
| 590 | Troxipide | other | 0.96 |
| 591 | Fluconazole | other | 1.03 |
| 592 | Ellagic | other | 1.08 |
| 593 | Fidaxomicin | other | 0.97 |
| 594 | Clodronate | other | 1.04 |
| 595 | Minocycline | other | 0.9 |
| 596 | Diminazene | other | 1.19 |
| 597 | Cinepazide | other | 1.11 |
| 598 | Sucralose | other | 0.99 |
| 599 | Praziquantel | other | 1.01 |
| 600 | Mevastatin | other | 0.64 |
| 601 | Suprofen | other | 1.01 |
| 602 | Doxycycline | other | 0.77 |
| 603 | Dirithromycin | other | 1.03 |
| 604 | Pemirolast | other | 0.63 |

| | | | |
|---|---|---|---|
| 605 | Ranolazine | other | 0.91 |
| 606 | Busulfan | other | 0.97 |
| 607 | Cisplatin | other | 0.83 |
| 608 | Dibenzepine | other | 0.78 |
| 609 | Cepharanthine | other | 0.83 |
| 610 | Phenacetin | other | 0.87 |
| 611 | Spectinomycin | other | 0.94 |
| 612 | Thonzonium | other | 1.05 |
| 613 | Thiostrepton | other | 0.97 |
| 614 | Camptothecin | other | 1.07 |
| 615 | Rolitetracycline | other | 0.96 |
| 616 | Rapamycin | other | 0.94 |
| 617 | Oxeladin | other | 0.99 |
| 618 | Carbadox | other | 1.04 |
| 619 | Piromidic | other | 1.06 |
| 620 | Deoxyarbutin | other | 0.9 |
| 621 | Monobenzone | other | 0.78 |
| 622 | Clindamycin | other | 0.99 |
| 623 | Pantothenic acid | other | 0.95 |
| 624 | Cephalomannine | other | 0.95 |
| 625 | Sarafloxacin | other | 0.95 |
| 626 | Pentoxifylline | other | 0.95 |
| 627 | Moxalactam | other | 1.05 |
| 628 | Camylofin | other | 0.9 |
| 629 | Benfotiamine | other | 0.65 |
| 630 | Methapyrilene | other | 0.43 |
| 631 | Clofazimine | other | 1.1 |
| 632 | Pentamidine | other | 1.13 |
| 633 | Cefaclor | other | 0.84 |
| 634 | Amoxicillin | other | 1.05 |
| 635 | Artemisinin | other | 0.92 |
| 636 | Telbivudine | other | 0.93 |
| 637 | Aniracetam | other | 0.79 |
| 638 | Catharanthine | other | 0.99 |
| 639 | Tranilast | other | 1.76 |
| 640 | Buflomedil | other | 1.14 |
| 641 | Lomefloxacin | other | 1.01 |
| 642 | Moroxydine | other | 0.88 |
| 643 | Ginkgolide | other | 1.05 |
| 644 | Metrizamide | other | 1.06 |
| 645 | Methylhydantoin | other | 1.1 |
| 646 | Voriconazole | other | 0.99 |
| 647 | Pridinol Methanesulfonate | other | 1.01 |
| 648 | Tioconazole | other | 0.96 |
| 649 | Penicillin | other | 0.91 |
| 650 | Flumequine | other | 1.08 |
| 651 | Atazanavir | other | 0.83 |

| | | | |
|---|---|---|---|
| 652 | Ofloxacin | other | 0.85 |
| 653 | Fenbendazole | other | 1.14 |
| 654 | Dextrose | other | 0.92 |
| 655 | Marbofloxacin | other | 0.82 |
| 656 | Pyrimethamine | other | 0.93 |
| 657 | Suxibuzone | other | 0.87 |
| 658 | Phthalylsulfacetamide | other | 0.92 |
| 659 | Phenothrin | other | 0.91 |
| 660 | Noscapine | other | 0.94 |
| 661 | Glafenine | other | 1.5 |
| 662 | Cinoxacin | other | 0.86 |
| 663 | aminohippurate Hydrate | other | 0.87 |
| 664 | Primaquine | other | 0.91 |
| 665 | Mepiroxol | other | 0.94 |
| 666 | Hemicholinium | other | 0.86 |
| 667 | Clofoctol | other | 1.09 |
| 668 | Cephapirin | other | 0.89 |
| 669 | Gluceptate | other | 0.89 |
| 670 | Butacaine | other | 0.71 |
| 671 | Auranofin | other | 2.62 |
| 672 | Aztreonam | other | 0.85 |
| 673 | Aminoacridine | other | 0.93 |
| 674 | Docetaxel | other | 0.66 |
| 675 | Alexidine | other | ND |
| 676 | Ethacridine | other | 0.99 |
| 677 | Potassium Iodide | other | 0.86 |
| 678 | Digoxigenin | other | 0.89 |
| 679 | Guanidine | other | 0.88 |
| 680 | Bentiromide | other | 0.85 |
| 681 | Fosfomycin | other | 0.64 |
| 682 | Difloxacin | other | 0.89 |
| 683 | Bekanamycin | other | 0.83 |
| 684 | Paclitaxel | other | 0.85 |
| 685 | Proadifen | other | 0.81 |
| 686 | ascorbate | other | 0.99 |
| 687 | Deoxycorticosterone | other | 0.81 |
| 688 | Cetrimonium Bromide | other | 1.89 |
| 689 | Norfloxacin | other | 0.97 |
| 690 | Bergapten | other | 0.98 |
| 691 | Bephenium | other | 0.79 |
| 692 | Diperodon | other | 0.77 |
| 693 | Isoxicam | other | 0.72 |
| 694 | Malotilate | other | 0.87 |
| 695 | Famprofazone | other | 0.82 |
| 696 | Piperacillin | other | 0.98 |
| 697 | Ifosfamide | other | 0.77 |
| 698 | Spiramycin | other | 0.75 |

| | | | |
|---|---|---|---|
| 699 | Phosphatidylcholine | other | 0.51 |
| 700 | Procodazole | other | 0.94 |
| 701 | Amorolfine | other | 0.85 |
| 702 | Chloramphenicol | other | 0.83 |
| 703 | Picrotoxinin | other | 0.72 |
| 704 | Pasiniazid | other | 0.96 |
| 705 | Sulbactam | other | 0.48 |
| 706 | Emetine | other | 0.67 |
| 707 | Streptozotocin | other | 0.66 |
| 708 | Mesalamine | other | 0.76 |
| 709 | Dimaprit | other | 0.81 |
| 710 | Dibenzothiophene | other | 0.95 |
| 711 | Colistimethate | other | 0.79 |
| 712 | Clorgyline | other | 0.86 |
| 713 | Clopamide | other | 0.56 |
| 714 | Hydrastinine | other | 0.99 |
| 715 | Clinafoxacin | other | 0.95 |
| 716 | Chromocarb | other | 0.93 |
| 717 | Ceftazidime | other | 0.83 |
| 718 | Nifenazone | other | 1.02 |
| 719 | Cephalexin | other | 0.93 |
| 720 | Meclocycline | other | 0.65 |
| 721 | Isosorbide | other | 0.86 |
| 722 | Azaguanine | other | 1.12 |
| 723 | Furaltadone | other | 0.82 |
| 724 | Ceftiofur | other | 1.03 |
| 725 | Resveratrol | other | 0.72 |
| 726 | Clindamycin | other | 0.71 |
| 727 | Levofloxacin | other | 0.75 |
| 728 | Riboflavin | other | ND |
| 729 | Dyclonine | other | 0.92 |
| 730 | Sorbitol | other | 0.92 |
| 731 | carnitine | other | 1.07 |
| 732 | Mannitol | other | 0.72 |
| 733 | Metronidazole | other | 0.92 |
| 734 | Menadione | other | 0.82 |
| 735 | Nalidixic acid | other | 0.97 |
| 736 | Cefprozil | other | 0.93 |
| 737 | Avobenzone | other | 1.03 |
| 738 | Artemether | other | 1.15 |
| 739 | Talc | other | 0.97 |
| 740 | Methoxsalen | other | 0.76 |
| 741 | Nicotinamide | other | 0.88 |
| 742 | Miconazole | other | 0.73 |
| 743 | Acetanilide | other | 0.79 |
| 744 | Sulfamethizole | other | 0.87 |
| 745 | Secnidazole | other | 0.84 |

| | | | |
|---|---|---|---|
| 746 | Famciclovir | other | 0.95 |
| 747 | Miconazole | other | 0.65 |
| 748 | Econazole nitrate | other | 0.73 |
| 749 | Adiphenine | other | 0.86 |
| 750 | Carnitine | other | 0.78 |
| 751 | Isoconazole | other | 0.75 |
| 752 | Scopine | other | 1.0 |
| 753 | Isoniazid | other | 0.69 |
| 754 | Clindamycin | other | 1.06 |
| 755 | Bisacodyl | other | 0.97 |
| 756 | Pramiracetam | other | 1.03 |
| 757 | Clarithromycin | other | 1.09 |
| 758 | Vidarabine | other | 0.78 |
| 759 | Aminolevulinic | other | 1.48 |
| 760 | Azacyclonol | other | 0.81 |
| 761 | Irsogladine | other | 1.23 |
| 762 | Amfebutamone | other | 1.18 |
| 763 | Alibendol | other | 1.12 |
| 764 | Mecarbinate | other | 1.14 |
| 765 | Moxifloxacin | other | 0.98 |
| 766 | Clindamycin | other | 0.83 |
| 767 | Rifaximin | other | 0.98 |
| 768 | Geniposidic | other | 0.93 |
| 769 | Sulfisoxazole | other | 1.15 |
| 770 | Genipin | other | 1.07 |
| 771 | Sulfamethoxazole | other | 0.99 |
| 772 | Geniposide | other | 1.0 |
| 773 | Pregnenolone | other | 0.99 |
| 774 | Paeoniflorin | other | 0.9 |
| 775 | Deacetylbaccatin | other | 0.85 |
| 776 | Cromoglycate | other | 0.94 |
| 777 | Sulbactam | other | 0.81 |
| 778 | Oxytetracycline | other | 1.07 |
| 779 | Nystatin | other | 0.88 |
| 780 | Crystal Violet | other | ND |
| 781 | Rebamipide | other | 0.77 |
| 782 | Acadesine | other | 0.82 |
| 783 | Fenticonazole | other | 0.63 |
| 784 | Azithromycin | other | 1.72 |
| 785 | Albendazole | other | 0.99 |
| 786 | Flunixin | other | 0.99 |
| 787 | Etidronate | other | 0.73 |
| 788 | Xylose | other | 0.71 |
| 789 | Raltegravir | other | 0.79 |
| 790 | Elvitegravir | other | 0.87 |
| 791 | Roxithromycin | other | 0.8 |
| 792 | orthovanadate | other | 0.91 |

| | | | |
|---|---|---|---|
| 793 | Ribavirin | other | 0.78 |
| 794 | Cycloserine | other | 0.68 |
| 795 | Liranaftate | other | 0.68 |
| 796 | Fudosteine | other | 0.7 |
| 797 | Quinine | other | 0.66 |
| 798 | Procarbazine | other | 0.79 |
| 799 | Licofelone | other | 0.78 |
| 800 | Bifonazole | other | 0.61 |
| 801 | Arbidol | other | 1.07 |
| 802 | Penicillamine | other | 0.83 |
| 803 | Probucol | other | 0.71 |
| 804 | Oxibendazole | other | 0.9 |
| 805 | Daidzein | other | 0.71 |
| 806 | Curcumin | other | 1.76 |
| 807 | Chloroxine | other | 1.16 |
| 808 | Vinpocetine | other | 0.67 |
| 809 | Lomustine | other | 0.69 |
| 810 | Novobiocin | other | 1.1 |
| 811 | Butoconazole | other | 0.7 |
| 812 | Valaciclovir | other | 0.5 |
| 813 | Oxfendazole | other | 1.25 |
| 814 | Ciclopirox | other | 0.81 |
| 815 | Methacycline | other | 0.8 |
| 816 | Lopinavir | other | 1.06 |
| 817 | Acipimox | other | 0.92 |
| 818 | Ciprofloxacin | other | 0.84 |
| 819 | Aciclovir | other | 0.81 |
| 820 | DAPT | other | 0.73 |
| 821 | Fleroxacin | other | 0.87 |
| 822 | Sulphadimethoxine | other | 0.83 |
| 823 | Rimantadine | other | 1.2 |
| 824 | Sparfloxacin | other | 1.01 |
| 825 | Primidone | other | 0.87 |
| 826 | Idoxuridine | other | 1.1 |
| 827 | Pivoxil | other | 0.96 |
| 828 | Protionamide | other | 0.71 |
| 829 | Itraconazole | other | 1.21 |
| 830 | Nefiracetam | other | 0.92 |
| 831 | Lincomycin | other | 0.85 |
| 832 | Chlormezanone | other | 1.0 |
| 833 | Cidofovir | other | 0.89 |
| 834 | Erdosteine | other | 0.96 |
| 835 | Suplatast | other | 0.84 |
| 836 | Tobramycin | other | 0.79 |
| 837 | Taurine | other | 1.29 |
| 838 | Sulfadoxine | other | 1.16 |
| 839 | Sitafloxacin | other | 0.77 |

| | | | |
|---|---|---|---|
| 840 | Ganciclovir | other | 0.83 |
| 841 | Trifluridine | other | 0.68 |
| 842 | Oseltamivir | other | 1.02 |
| 843 | Verteporfin | other | ND |
| 844 | Valganciclovir | other | 0.88 |
| 845 | Cyclandelate | other | 1.01 |
| 846 | Antipyrine | other | 0.84 |
| 847 | Sasapyrine | other | 0.96 |
| 848 | Enoxacin | other | 0.89 |
| 849 | Salicylanilide | other | 0.87 |
| 850 | Ampicillin | other | 0.91 |
| 851 | Domiphen | other | 1.22 |
| 852 | Abacavir | other | 0.68 |
| 853 | Sulfacetamide | other | 0.9 |
| 854 | Amoxicillin | other | 0.91 |
| 855 | Linezolid | other | 0.93 |
| 856 | Rifapentine | other | 1.2 |
| 857 | L-Arginine | other | 1.02 |
| 858 | Amprenavir | other | 0.99 |
| 859 | Zanamivir | other | 0.59 |
| 860 | Mequinol | other | 0.64 |
| 861 | Albendazole | other | 1.12 |
| 862 | L-Thyroxine | other | 1.11 |
| 863 | Carbazochrome | other | 0.93 |
| 864 | Flucytosine | other | 0.62 |
| 865 | Hygromycin | other | 0.71 |
| 866 | Aminosalicylate | other | 0.7 |
| 867 | Decamethonium | other | 0.87 |
| 868 | Paromomycin Sulfate | other | 0.88 |
| 869 | Tylosin tartrate | other | 0.98 |
| 870 | Nifuroxazide | other | 0.85 |
| 871 | Posaconazole | other | 1.21 |
| 872 | Sertaconazole | other | 0.62 |
| 873 | Cinchophen | other | 0.96 |
| 874 | Chlorquinaldol | other | 0.97 |
| 875 | Azlocillin | other | 0.88 |
| 876 | Florfenicol | other | 0.73 |
| 877 | Tolperisone | other | 0.85 |
| 878 | Octopamine | other | 0.72 |
| 879 | Vinblastine | other | 1.13 |
| 880 | Aminothiazole | other | 0.91 |
| 881 | Bemegride | other | 1.05 |
| 882 | Carboplatin | other | 0.98 |
| 883 | Erythromycin | other | 1.04 |
| 884 | Amphotericin | other | 0.96 |
| 885 | Niclosamide | other | 0.91 |
| 886 | Fenspiride | other | 1.04 |

| | | | |
|---|---|---|---|
| 887 | Betamipron | other | 1.0 |
| 888 | PMSF | other | 1.0 |
| 889 | Teicoplanin | other | 0.88 |
| 890 | Cabazitaxel | other | 1.0 |
| 891 | Tenofovir | other | 0.83 |
| 892 | Tenofovir | other | 1.02 |
| 893 | Glutamine | other | 0.85 |
| 894 | Ciclopirox | other | 0.78 |
| 895 | Tigecycline | other | 0.88 |
| 896 | Gadodiamide | other | 0.95 |
| 897 | Broxyquinoline | other | 1.44 |
| 898 | Carbenicillin | other | 0.98 |
| 899 | Chenodeoxycholic | other | 0.74 |
| 900 | Entecavir Hydrate | other | 0.86 |
| 901 | Stavudine | other | 0.98 |
| 902 | Nefopam | other | 0.86 |
| 903 | Hexadecanol | other | 0.97 |
| 904 | Naftifine | other | 1.0 |
| 905 | Sulfadiazine | other | 0.76 |
| 906 | Dehydroepiandrosterone | other | 1.04 |
| 907 | Idebenone | other | 1.2 |
| 908 | Erythromycin | other | 1.06 |
| 909 | Retapamulin | other | 1.0 |
| 910 | Trimethoprim | other | 1.01 |
| 911 | Oxytetracycline | other | 0.99 |
| 912 | Ranolazine | other | 0.81 |
| 913 | Cytarabine | other | 1.14 |
| 914 | Ronidazole | other | 1.13 |
| 915 | Cetylpyridinium | other | 1.41 |
| 916 | Carotene | other | 0.87 |
| 917 | Coumarin | other | 0.97 |
| 918 | Biotin | other | 1.0 |
| 919 | Pefloxacin Mesylate | other | 0.96 |
| 920 | Methenamine | other | 1.03 |
| 921 | Cyromazine | other | 0.96 |
| 922 | Benzethonium | other | 1.82 |
| 923 | Cefditoren | other | 0.84 |
| 924 | Sulfamerazine | other | 1.02 |
| 925 | Rifampin | other | 0.84 |
| 926 | Clorsulon | other | 1.1 |
| 927 | Sulfamethazine | other | 0.96 |
| 928 | Nitrofural | other | 0.69 |
| 929 | Sulfaguanidine | other | 1.08 |
| 930 | Trometamol | other | 1.0 |
| 931 | Tianeptine | other | 0.82 |
| 932 | Deferiprone | other | 0.8 |
| 933 | Netilmicin | other | 0.99 |

| | | | |
|---|---|---|---|
| 934 | Pyrazinamide | other | 0.93 |
| 935 | Tinidazole | other | 0.8 |
| 936 | Peramivir | other | 1.23 |
| 937 | Climbazole | other | 0.92 |
| 938 | Dequalinium | other | 1.29 |
| 939 | Tioxolone | other | 1.08 |
| 940 | Mezlocillin | other | 0.95 |
| 941 | Arecoline | other | 1.0 |
| 942 | Butenafine | other | 1.16 |

Comounds in plates with the Z'-factor below the threshold.

| # | **Name** | **Target** |
|---|---|---|
| 1 | Silodosin | membrane receptor |
| 2 | Pizotifen | membrane receptor |
| 3 | Fumarate | membrane receptor |
| 4 | Clorprenaline | membrane receptor |
| 5 | Naloxone | membrane receptor |
| 6 | Brompheniramine | membrane receptor |
| 7 | Mianserin | membrane receptor |
| 8 | Telmisartan | membrane receptor |
| 9 | Mosapride | membrane receptor |
| 10 | Risperidone | membrane receptor |
| 11 | Methylsulfate | membrane receptor |
| 12 | Trazodone | membrane receptor |
| 13 | Homatropine | membrane receptor |
| 14 | Melatonin | membrane receptor |
| 15 | Bisoprolol fumarate | membrane receptor |
| 16 | Olmesartan | membrane receptor |
| 17 | Imiquimod | membrane receptor |
| 18 | Hydroxyzine | membrane receptor |
| 19 | Indacaterol | membrane receptor |
| 20 | Dexmedetomidine | membrane receptor |
| 21 | Rizatriptan | membrane receptor |
| 22 | Phenylephrine | membrane receptor |
| 23 | Candesartan | membrane receptor |
| 24 | Oxybutynin | membrane receptor |
| 25 | Isoprenaline | membrane receptor |
| 26 | Lafutidine | membrane receptor |
| 27 | Aclidinium | membrane receptor |
| 28 | baclofen | membrane receptor |
| 29 | Paliperidone | membrane receptor |
| 30 | Tetrahydrozoline | membrane receptor |
| 31 | Levodopa | membrane receptor |
| 32 | Roxatidine | membrane receptor |
| 33 | Terazosin | membrane receptor |
| 34 | Xylometazoline | membrane receptor |
| 35 | Loratadine | membrane receptor |

| | | |
|---|---|---|
| 36 | Fludarabine | enzyme |
| 37 | Doxorubicin | enzyme |
| 38 | Indomethacin | enzyme |
| 39 | Fluorouracil | enzyme |
| 40 | Methotrexate | enzyme |
| 41 | Clofarabine | enzyme |
| 42 | Methazolamide | enzyme |
| 43 | Idarubicin | enzyme |
| 44 | Apixaban | enzyme |
| 45 | Quinapril | enzyme |
| 46 | Pyridostigmine | enzyme |
| 47 | Lovastatin | enzyme |
| 48 | Tolcapone | enzyme |
| 49 | Decitabine | enzyme |
| 50 | Epirubicin | enzyme |
| 51 | Hydralazine | enzyme |
| 52 | Dexrazoxane | enzyme |
| 53 | Etoposide | enzyme |
| 54 | Ubenimex | enzyme |
| 55 | Flavoxate | enzyme |
| 56 | Dorzolamide | enzyme |
| 57 | Mitotane | enzyme |
| 58 | Dutasteride | enzyme |
| 59 | Gemcitabine | enzyme |
| 60 | Fenoprofen | enzyme |
| 61 | Piroxicam | enzyme |
| 62 | Miglitol | enzyme |
| 63 | Omeprazole | enzyme |
| 64 | Carprofen | enzyme |
| 65 | Mizoribine | enzyme |
| 66 | Ketoconazole | enzyme |
| 67 | Lansoprazole | enzyme |
| 68 | Meloxicam | enzyme |
| 69 | Riluzole | ion channel |
| 70 | Granisetron | ion channel |
| 71 | Penfluridol | ion channel |
| 72 | Oxcarbazepine | ion channel |
| 73 | Procaine | ion channel |
| 74 | Prilocaine | ion channel |
| 75 | Dibucaine | ion channel |
| 76 | Lidocaine | ion channel |
| 77 | Tetracaine | ion channel |
| 78 | Amiloride | ion channel |
| 79 | Ondansetron | ion channel |
| 80 | Rosiglitazone | nuclear receptor |
| 81 | Beclomethasone | nuclear receptor |
| 82 | Medroxyprogesterone | nuclear receptor |

| | | |
|---|---|---|
| 83 | Prednisolone | nuclear receptor |
| 84 | Clomifene | nuclear receptor |
| 85 | Cortisone | nuclear receptor |
| 86 | Prednisolone | nuclear receptor |
| 87 | Ulipristal | nuclear receptor |
| 88 | Bezafibrate | nuclear receptor |
| 89 | Norethindrone | nuclear receptor |
| 90 | Vitamin C | nuclear receptor |
| 91 | Pioglitazone | nuclear receptor |
| 92 | Gestodene | nuclear receptor |
| 93 | Drospirenone | nuclear receptor |
| 94 | Isotretinoin | nuclear receptor |
| 95 | Gemfibrozil | nuclear receptor |
| 96 | Estradiol | nuclear receptor |
| 97 | Levonorgestrel | nuclear receptor |
| 98 | Eplerenone | nuclear receptor |
| 99 | Methylprednisolone | nuclear receptor |
| 100 | Clobetasol | nuclear receptor |
| 101 | Progesterone | nuclear receptor |
| 102 | Prednisone | nuclear receptor |
| 103 | Acitretin | nuclear receptor |
| 104 | Raloxifene | nuclear receptor |
| 105 | Levetiracetam | transporter |
| 106 | Hydrochlorothiazide | transporter |
| 107 | Minoxidil | transporter |
| 108 | Furosemide | transporter |
| 109 | Glyburide | transporter |
| 110 | Metolazone | transporter |
| 111 | Reserpine | transporter |
| 112 | Glimepiride | transporter |
| 113 | Pidotimod | other |
| 114 | Ornidazole | other |
| 115 | Chloroquine | other |
| 116 | Sulfathiazole | other |
| 117 | Chlorzoxazone | other |
| 118 | Caspofungin | other |
| 119 | Dropropizine | other |
| 120 | Flubendazole | other |
| 121 | Pyridoxine | other |
| 122 | D-Phenylalanine | other |
| 123 | Eprazinone | other |
| 124 | Chlortetracycline | other |
| 125 | Thiamphenicol | other |
| 126 | Ethamsylate | other |
| 127 | Vitamin D2 | other |
| 128 | Olsalazine | other |
| 129 | levofolinate | other |

| | | |
|---|---|---|
| 130 | Zidovudine | other |
| 131 | Nafcillin | other |
| 132 | Ampicillin | other |
| 133 | Azithromycin | other |
| 134 | Amprolium | other |
| 135 | Toltrazuril | other |
| 136 | Bacitracin | other |
| 137 | Acetylcysteine | other |
| 138 | Orbifloxacin | other |
| 139 | Doxapram | other |
| 140 | Moguisteine | other |
| 141 | Atovaquone | other |
| 142 | Nadifloxacin | other |
| 143 | Creatinine | other |
| 144 | Decoquinate | other |
| 145 | Rifabutin | other |
| 146 | Nevirapine | other |
| 147 | Sulfapyridine | other |
| 148 | Amikacin | other |
| 149 | Cefoperazone | other |
| 150 | Clafen | other |
| 151 | Altretamine | other |
| 152 | Terbinafine | other |
| 153 | Doripenem | other |
| 154 | Azathioprine | other |
| 155 | Daptomycin | other |
| 156 | Zalcitabine | other |
| 157 | Cefoselis | other |
| 158 | Adefovir | other |
| 159 | Biapenem | other |
| 160 | Bleomycin | other |
| 161 | Nelarabine | other |
| 162 | Clotrimazole | other |
| 163 | Bendamustine | other |
| 164 | Cefdinir | other |
| 165 | Deferasirox | other |
| 166 | Ivermectin | other |
| 167 | Lamivudine | other |
| 168 | Sulfameter | other |
| 169 | Darunavir | other |
| 170 | Dacarbazine | other |
| 171 | Oxacillin | other |
| 172 | Streptomycin | other |
| 173 | Probenecid | other |
| 174 | Neomycin | other |
| 175 | Picosulfate | other |
| 176 | Cloxacillin | other |

| | | |
|---|---|---|
| 177 | Colistin | other |
| 178 | Meropenem | other |
| 179 | Balofloxacin | other |
| 180 | Tiopronin | other |
| 181 | Bromhexine | other |
| 182 | Tolnaftate | other |
| 183 | Cyclophosphamide | other |
| 184 | Docosanol | other |
| 185 | Guaifenesin | other |
| 186 | Thiabendazole | other |
| 187 | Vancomycin | other |
| 188 | Nafamostat | other |
| 189 | Methocarbamol | other |
| 190 | Mesna | other |
| 191 | Terbinafine | other |
| 192 | Tetracycline | other |
